# Fructose consumption in pregnancy and associations with maternal and offspring hepatic and whole-body adiposity: a scoping review

**DOI:** 10.1101/2024.07.02.600389

**Authors:** Grace Zhao, Sarah Chondon, Clint Gray, Sheridan Gentili, Meagan Stanley, Timothy RH Regnault

## Abstract

**Background:** Fructose is a major component in the Western diet, and its increased intake has been linked to adverse metabolic health, including impaired hepatic function and increased adiposity. The early life period, including preconceptionally, pregnancy and the newborn period, are critical periods in determining later metabolic health. However, the impact of excess fructose intake during this time on maternal, fetal, and offspring hepatic and whole-body adiposity, are ill defined.

**Objectives:** To understand the effects of maternal fructose consumption pre- and during pregnancy on maternal, fetal and offspring hepatic and whole-body adiposity.

**Methods:** A systematic search of MEDLINE, EMBASE, and CENTRAL was performed up to August 23, 2022, to identify studies that focused on maternal fructose consumption pre- and during pregnancy on hepatic and whole-body adiposity in the mother, fetus, and offspring. Citations, abstracts, and full texts were screened in duplicate. Hepatic adiposity was defined as elevated hepatic triglycerides or overall hepatic fat accumulation. Whole-body adiposity was defined as increased adipose tissue or adipocyte hypertrophy.

**Results:** After screening 2334 citations, 33 experimental studies reporting maternal fructose consumption pre- and during pregnancy in rodents were included. Prenatal fructose exposure was associated with maternal (9 out of 12) and offspring (6 out of 10) whole-body adiposity. A high proportion of studies (13 out of 14) supported the association between fructose during pregnancy and increased maternal hepatic adiposity. Fetal hepatic adiposity and elevated expression of hepatic lipogenic proteins were noted in four studies. Offspring hepatic adiposity was supported in 14 of the 17 articles that discussed hepatic results, with five studies demonstrating more severe effects in female offspring.

**Conclusions:** Fructose consumption during pregnancy in rodent models is associated with maternal, fetal, and offspring hepatic, whole-body adiposity and underlying sex-specific effects. There are no human fructose studies and its effects in the early life period.

**Registration number:** H8F26 on Open Science Framework

## INTRODUCTION

Fructose has become an increasingly significant component of the carbohydrate content in diets due to its high sweetness value.^1-3^ Excessive fructose consumption is associated with non-alcoholic fatty liver disease (NAFLD), obesity, weight gain, and along with hyperuricemia, inflammation and insulin resistance,^4-6^ which are key components of metabolic syndrome.^7-10^ Fructose consumption is considered one of the primary causes of metabolic syndrome in adults.^11^ A healthy diet during pregnancy is essential to support the optimal growth and development of the fetus. However, data show that fewer pregnant women are meeting current diet recommendations.^12^ Studies highlight that these dietary changes are associated with the increased consumption of low nutrient-dense foods, including fructose-sweetened foods, which now appear to be the major contributors to the total energy intake in women during pregnancy.^13-15^

Metabolically, intestinal fructose uptake is mediated by the glucose transporter GLUT5 (Slc2a5) in the apical membrane and GLUT2 (Slc2a2) in the basolateral membrane.^16^ Within the intestinal epithelial enterocytes, fructose is phosphorylated by KHK and converted to glucose, lactate, glycerate, and other organic acids, in addition to modulating the composition of gut microbiota and transcription of GLUT5 for further uptake.^17, 18^ However, the majority of dietary fructose exits the enterocyte via GLUT2 to enter the portal vein and into the liver.^18^ Here, fructose is extracted by hepatocytes, likely through GLUT2 transporter and phosphorylated by KHK to F1P, and then its metabolites act as signaling molecules to activate metabolic transcriptional programs associated with glucose production, lipogenesis, and glycogen synthesis.^16^ Importantly, hepatic fructose metabolism is considered to lead to cellular depletion of ATP^19, 20^ and also activates transcription factors including ChREBP and SREBP1c, which regulate enzymes involved in fructolysis, glycolysis, glucose production, lipogenesis, and VLDL packaging and export,^21, 22^ processes that underly hepatic *de novo* lipogenesis.^23^ This possibly leads to the development of NALFD, now a public health issue.^24^

In pregnancy, the hexose transporters GLUT8 (Scl2a8) and GLUT9 (Scl2a9), which transport fructose, are expressed in human placentae,^25, 26^ in addition to the presence of KHK in trophoblastic tissues.^27^ Fructose is an important placental product^28-30^ from glucose, and a substrate for normal placental development, playing a key role during the relatively hypoxic early placentation period, to maintain ATP concentrations and cellular redox potentials.^30, 31^ During the early period of pregnancy, fructose is perhaps preferred over glucose in relatively hypoxic tissues. Fructose can be endogenously produced from glucose in some pathological conditions. Furthermore, fructose production normally decreases significantly following the establishment of maternal-fetal circulation following placentation^31^. Negative impacts of excessive fructose consumption are demonstrated in both non-pregnant human and animal studies, ^11, 32^, and also in pregnancy. In pregnant mice and rat models, excessive fructose consumption is associated with placental insufficiency, elevated uric acid synthesis and abnormal fetal growth.^33^ In humans, maternal serum fructose levels are correlated with a risk of gestational diabetes,^34-36^ pre-eclampsia,^37^ and placental uric acid levels, suggesting similar uric acid pathways, as exhibited in rodent models, might also be functioning in human placenta.^38^ Further animal studies have shown fructose to be a strong independent risk factor for preeclampsia, possibly through uric acid-associated endothelial dysfunction factors,^31^ and H2S/angiogenic mechanisms,^39^ though further work is ongoing to better understand fructose’s possible contributions to preeclampsia and other adverse pregnancy outcome situations. Additionally, studies show that exposure to certain flavors through amniotic fluid and breast milk has an influence on the development of taste preferences^40, 41^ and links with increased whole-body adiposity and NAFLD are emerging.^42, 43^ While studies highlight the adverse impact of high-fructose diets on metabolic health in the non-pregnant state, the understanding of fructose’s impact in the periconceptional and pregnant state, specifically on human maternal and offspring body adiposity and hepatic fat content is still developing.

Animal studies have shown that consumption of excessive fructose during pregnancy has had adverse effects on fetal development and in later life.^44, 45^ However, there is a paucity of human data on this topic. Further, the short and long-term impacts of elevated fructose consumption on hepatic and whole-body adiposity, early markers of later life metabolic health, in the early life period still needed to be elucidated. We therefore conducted a scoping review of the current literature to examine the relationship between fructose consumption during pre-pregnancy, pregnancy, and lactation, and its effects on maternal and offspring hepatic and whole-body adiposity.

## METHODS

A scoping review was conducted with guidance from the scoping review methodological framework developed by Arksey and O’Malley,^46^ the Synthesis Without Meta-Analysis guideline,^47^ and the JBI methodology for scoping review.^48^ The protocol was prospectively registered with Open Science Framework (identifier H8F26).

### Search strategy

The search strategy was developed in consultation with a health sciences information specialist (MS) and searched MEDLINE (Ovid), EMBASE (Ovid), Cochrane Central Register of Controlled Trials (CENTRAL) from inception to August 23, 2022, for human and animal studies with no language restrictions. The full search strategy can be found in Appendix A. Bibliographies of included studies and relevant reviews were reviewed to identify additional studies.

### Eligibility Criteria

Study eligibility was defined using the following PECO (population, exposure, comparison group, outcome) criteria: 1) pregnant dams or women; 2) maternal fructose consumption (pre-pregnancy, during pregnancy to throughout lactation period); 3) included a control group without maternal fructose consumption; and 4) reported hepatic adiposity (liver weight, hepatic triglycerides, histological hepatic lipid droplets) or whole-body adiposity (adipose tissue, weight gain). Studies focusing on obstetrical outcomes, and abstracts, conference proceedings, letters to the editor, and literature reviews were excluded.

### Study selection

All identified citations were uploaded into Covidence, and duplicates removed. Following a pilot test, abstracts and full texts were assessed for eligibility in duplicate by independent reviewers (two of GZ, SC, TRHR, GC, SG). Disagreements on study eligibility were discussed with the study team and resolved by consensus. Notably, the eligibility criteria were modified at the full text screen such that studies that used fructose in combination with other diets instead of in isolation were excluded.

### Data Extraction

Data extraction was performed by GZ and SC and verified by TRHR using a data extraction tool developed by the reviewers. We conducted a pilot extraction of five articles to address any concerns with the article selection criteria and the extraction tool. For included articles, we extracted study characteristics (*i.e*. year, region), population characteristics (i.e. animal species, follow-up timeline), exposure characteristics (*i.e*. fructose concentration and timing of administration), and hepatic fat accumulation or whole-body adiposity characteristics (maternal, fetal, placental, or offspring), and potential mechanisms of adiposity. Fructose exposure was categorized as high (>30% wt:vol or wt:wt) or low (<30% wt:vol or wt:wt) to simplify the variation of fructose concentrations. The threshold of 30% was chosen as studies have shown that levels above this exceeds the average human intake and is not physiological.^49-51^ We categorized hepatic fat accumulation as elevated hepatic triglyceride content and macro- and micro-vesicular fatty infiltration in hepatocytes. We considered whole-body adiposity as an increase in adipose tissue (retroperitoneal, visceral, white) or adipocyte hypertrophy. Data for hepatic fat accumulation and whole-body adiposity measures is presented in a narrative format with tables and figures.

## RESULTS

### Study characteristics

The systematic search yielded 2,334 citations after duplicates were removed, of which 33 met the inclusion criteria (**Figure 1** – PRISMA). The included 33 records were published between 1990-2022, with 21 (64%) published between 2000-2019 (**Table 1**). Individual study characteristics are presented in **Table 2**. All included studies were experimental studies on rodents and no human reports met the inclusion criteria. Most included studies were conducted in North America (n = 8, 24%) and Oceania (n = 8, 24%). While the timing and duration of fructose exposure differed between studies, the two most common timings were during pregnancy only (n=12, 36%) and during pregnancy and lactation (n=11, 33%). High-dose fructose solution (>30% wt:vol or wt:wt) was used in six studies^45, 52-56^ and low-dose fructose solution (<30% wt:vol or wt:wt) was used in 21studies.^42, 57-76^ The time points for data collection ranged from day 18 of gestation to one year postnatal. Data collection timepoints varied between studies and are described in Table 2 and visualized in **Figure 2**. Of interest, six studies only studied male offspring,^42, 53, 55, 59, 65, 75^ and two studies only studied female offspring.^68, 76^

**Fig. 1.**
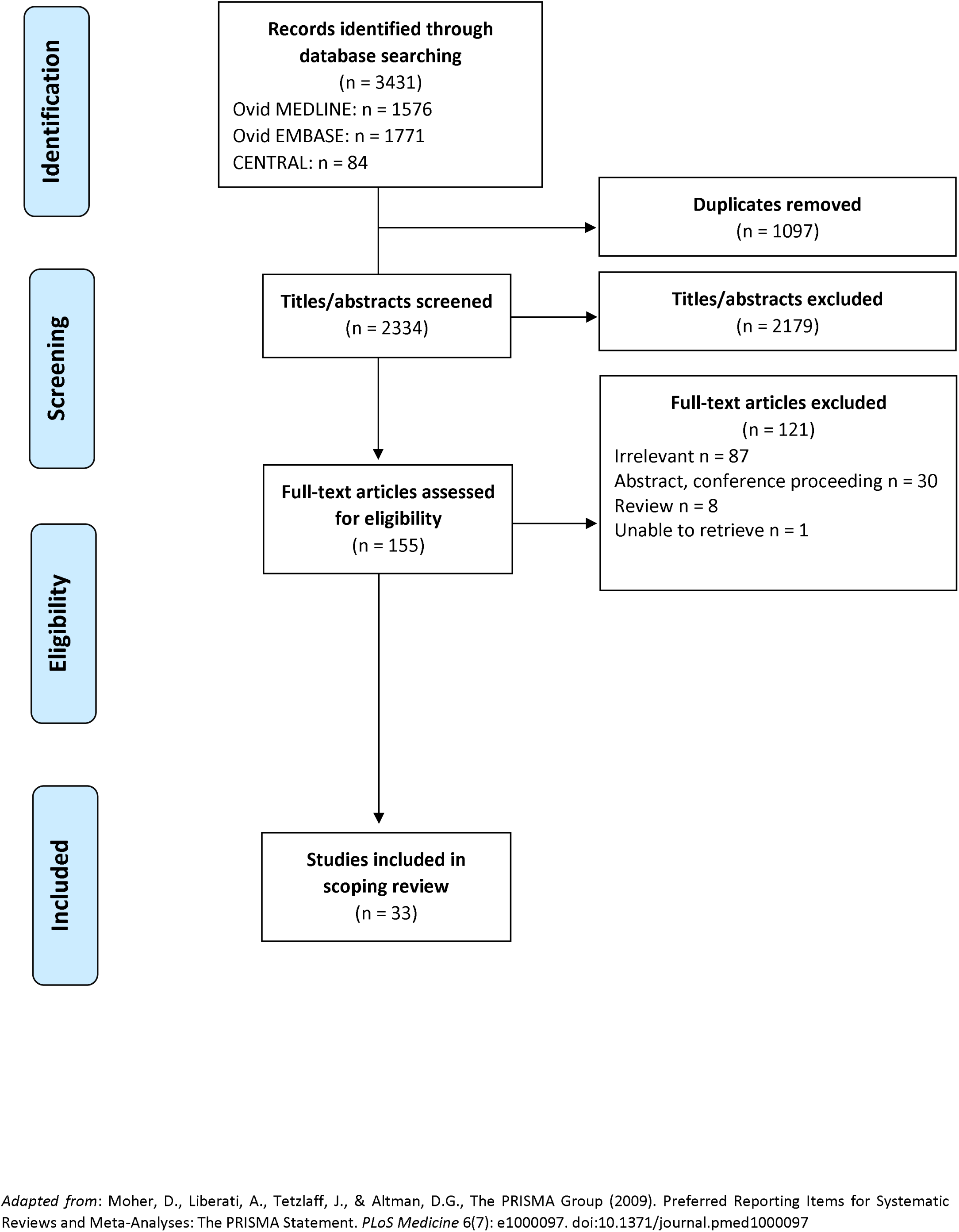
PRISMA flow diagram of literature search and selection process.

**Fig. 2.**
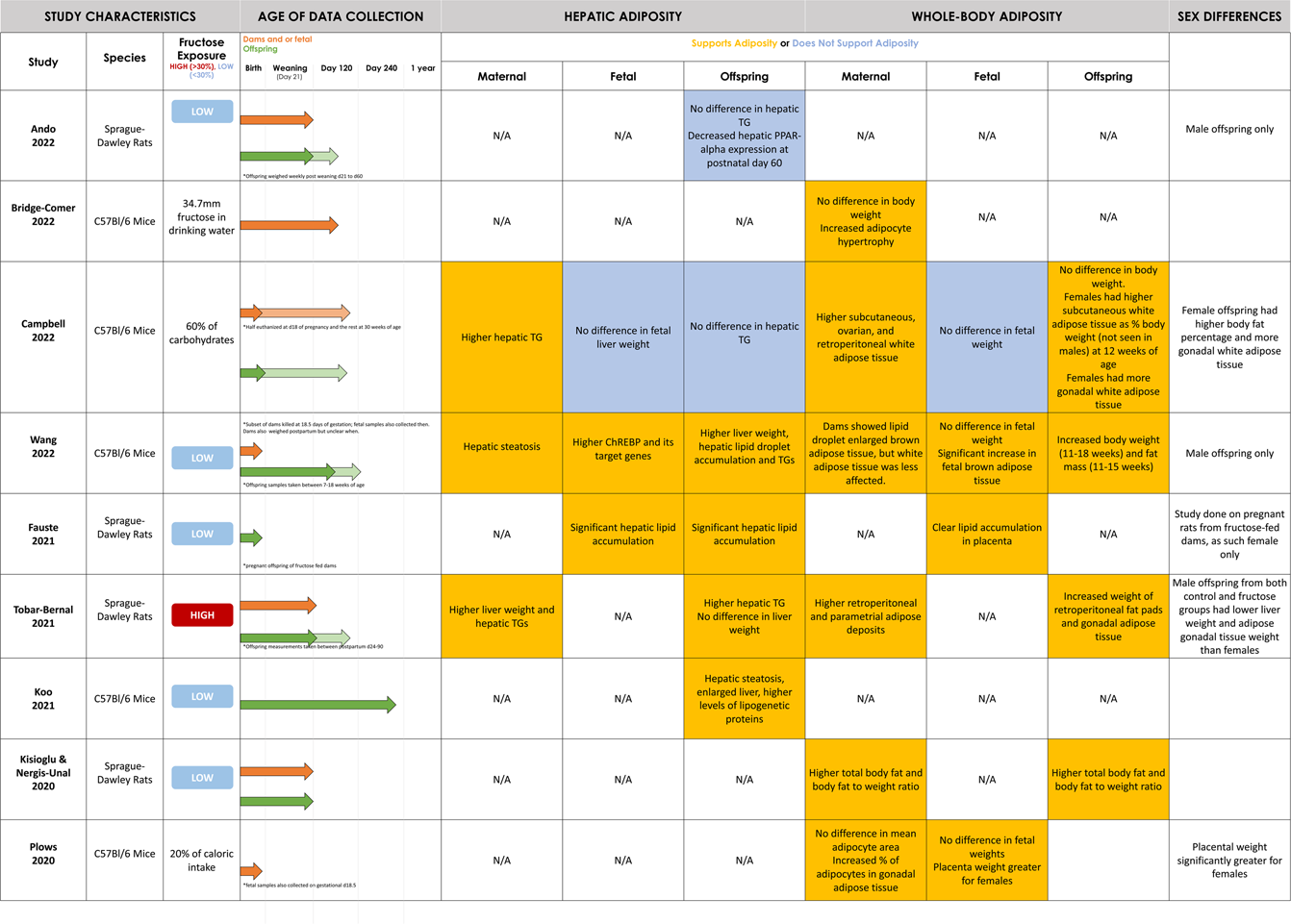
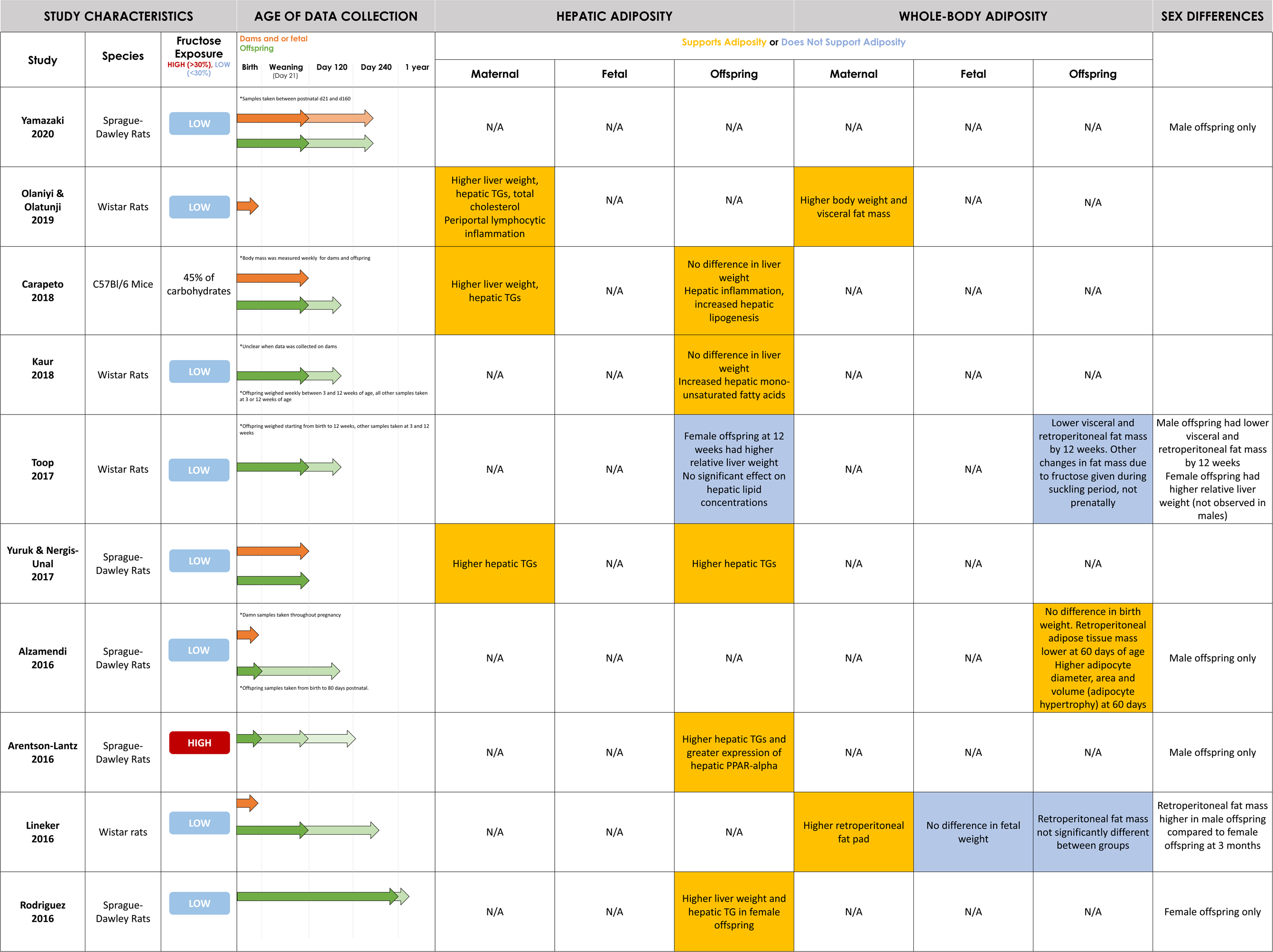
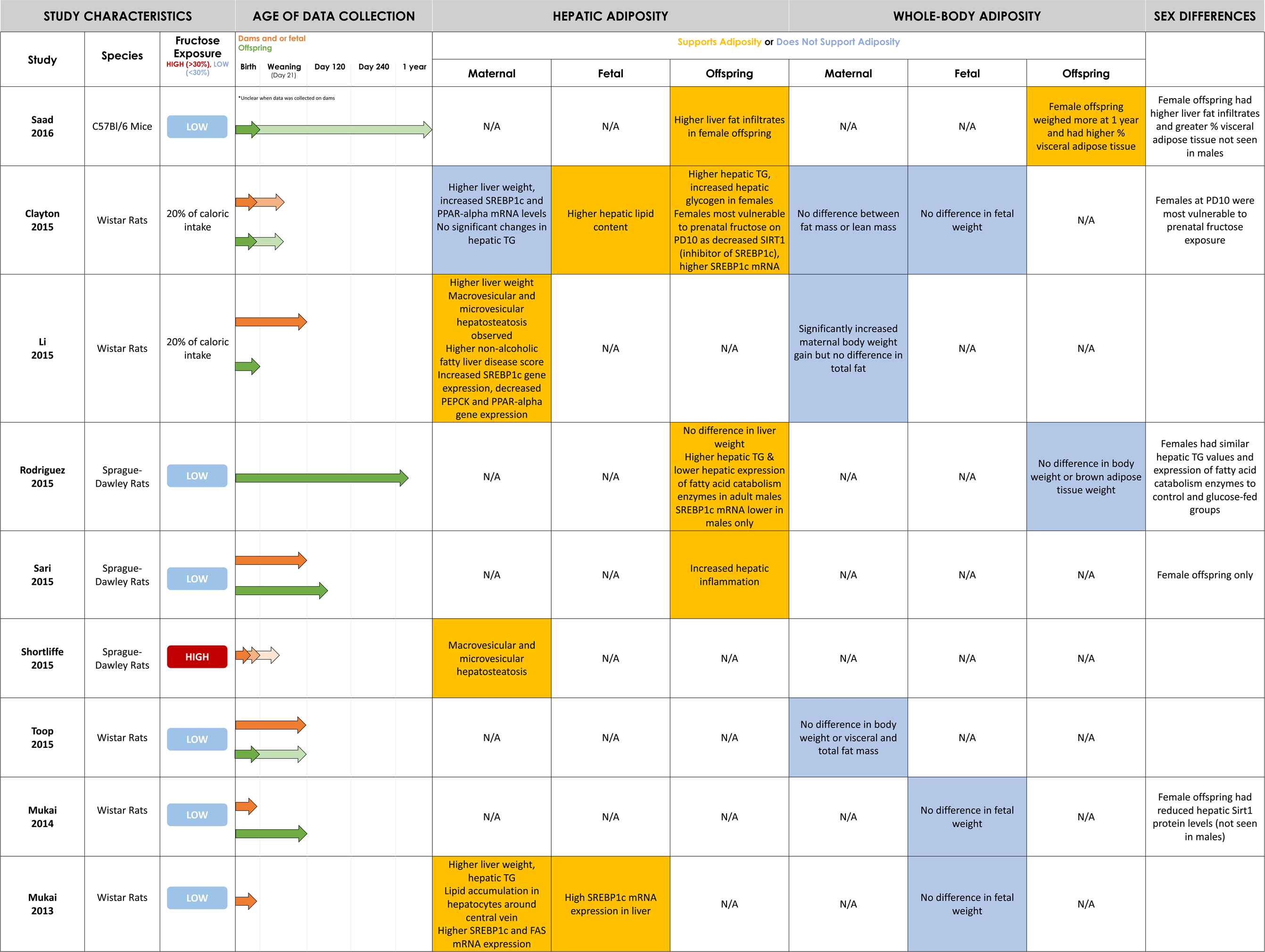
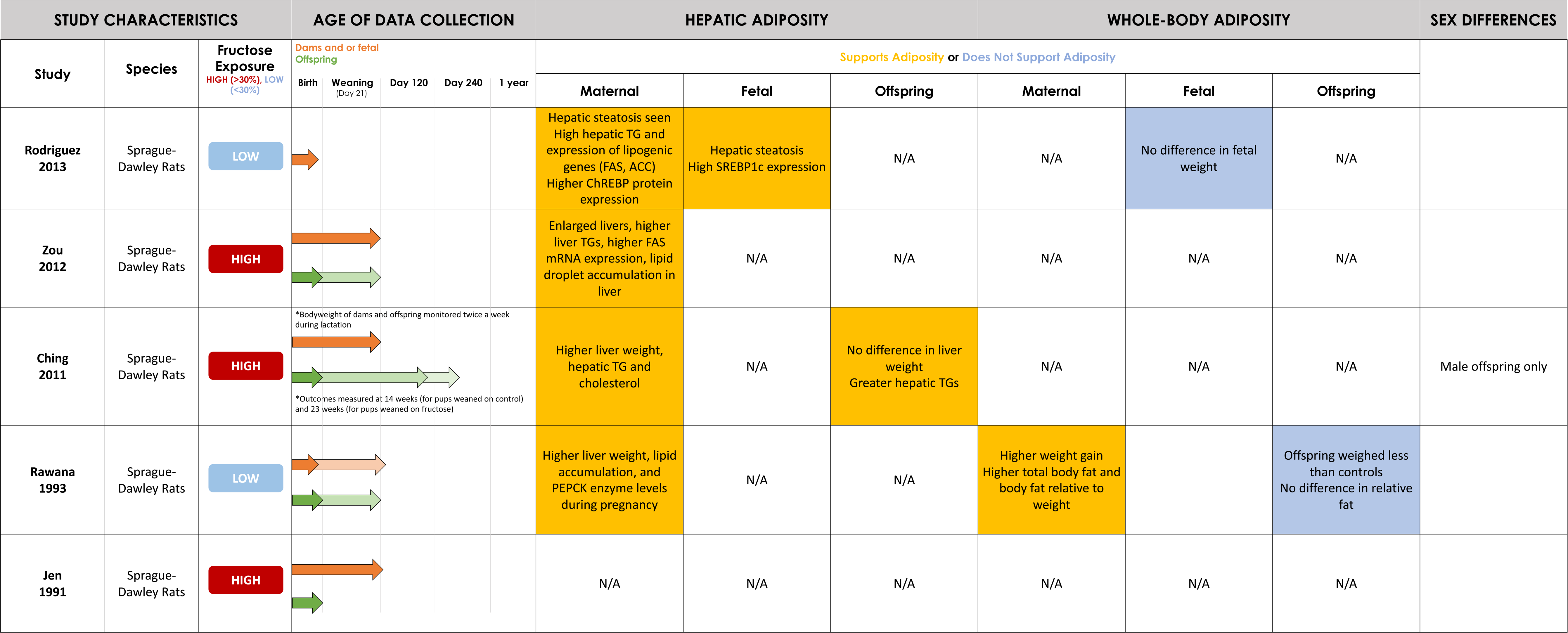
Summary of included study characteristics and hepatic and whole-body adiposity findings.

**Table 1:** Overview of selected article characteristics (n=33 total)

| Characteristic | Output |
| --- | --- |
| Year | • 1990-1999 (6%, n=2)<br>• 2000-2019 (64%, n=21)<br>• 2020-2022 (30%, n=10) |
| Geography | • Oceania (24%, n=8)<br>• North America (24%, n=8)<br>• Asia (21%, n=7)<br>• Europe (12%, n=4)<br>• South America (6%, n=2)<br>• Africa (3%, n=1)<br>• Other: Turkey (9%, n=3) |
| Species | • Sprague-Dawley Rats (52%, n=17)<br>• C57BL/6J Mice (21%, n=7)<br>• Wistar Rats (27%, n=9) |
| Fructose Exposure Timing | • Dams Pregnancy Only (36%, n=12)<br>• Dams Pregnancy and Lactation (33%, n=11)<br>• Dams Pregnancy and Lactation, and Offspring Post-Weaning (3%, n=1)<br>• Dams Pre-Pregnancy and Pregnancy (3%, n=1)<br>• Dams Pre-Pregnancy, Pregnancy and Lactation (15%, n=5)<br>• Dams Pre-Pregnancy, Pregnancy and Lactation, and Offspring Post-Weaning (3%, n=1)<br>• Dams Pregnancy and Offspring Post-Weaning (3%, n=1)<br>• Dams Pregnancy and Pregnant Offspring (3%, n=1) |
| % Fructose | • 10-29% wt/vol solution (45% n=15)<br>• 100 g/L (9%, n=3)<br>• 10-29% solution (9%, n=3)<br>• 30-65% solution (12%, n= 4)<br>• 50-60% wt:wt (6%, n=2)<br>• Other:<br>• 20% of caloric intake (9%, n=3)<br>• 45-60% of carbohydrate intake (6%, n=2)<br>• 34.77mm (3%, n=1) |
| % Studies Supporting Maternal Hepatic Adiposity (of 14 relevant articles) | • 93% (n=13) |
| % Studies Supporting Offspring Hepatic Adiposity (of 17 relevant articles) | • 82% (n=14) |
| % Studies Supporting Maternal Whole-Body Adiposity (of 12 relevant articles) | • 75% (n=9) |
| % Studies Supporting Offspring Whole-Body Adiposity (of 10 relevant articles) | • 60% (n=6) |

**Table 2:**
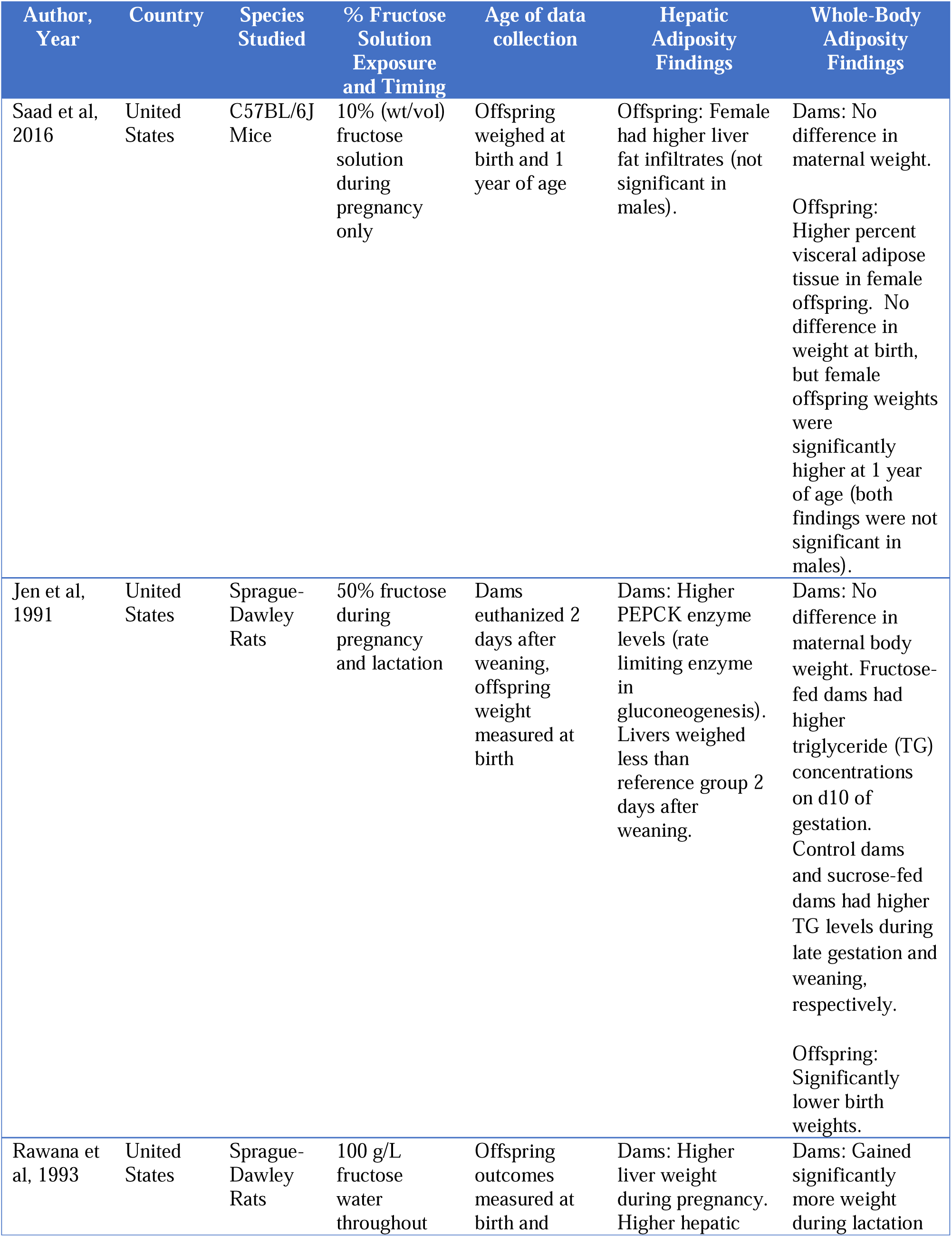
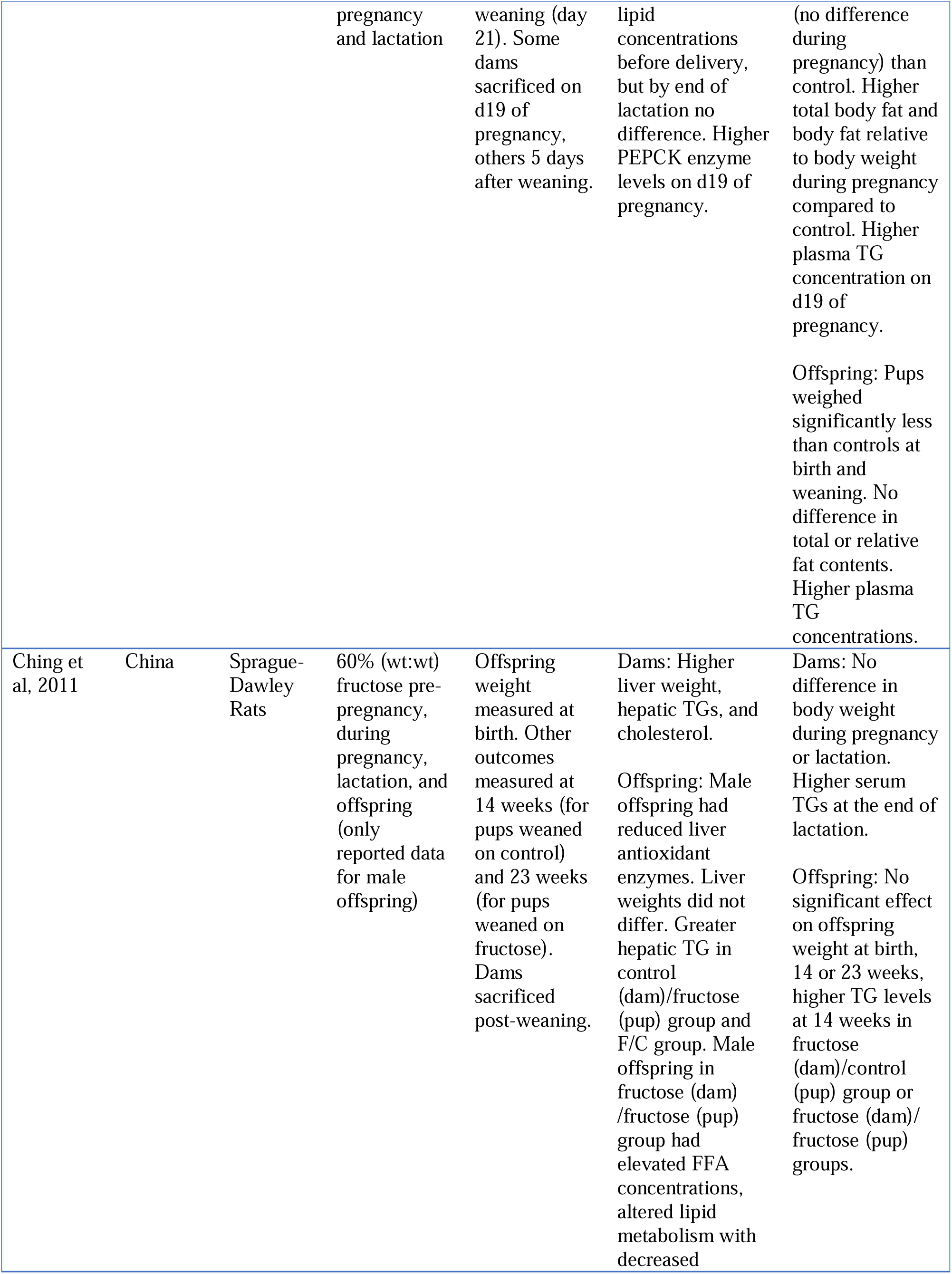
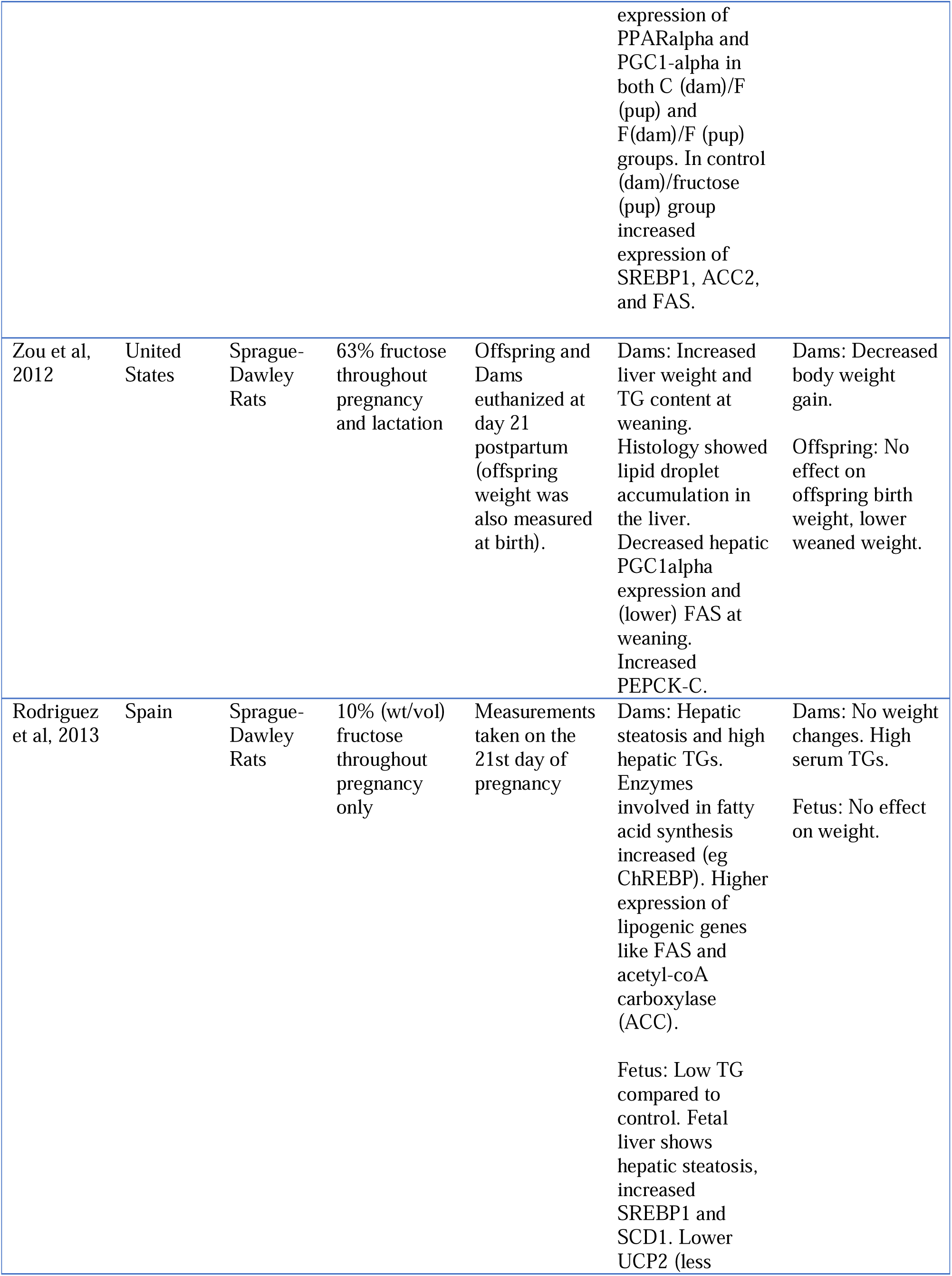
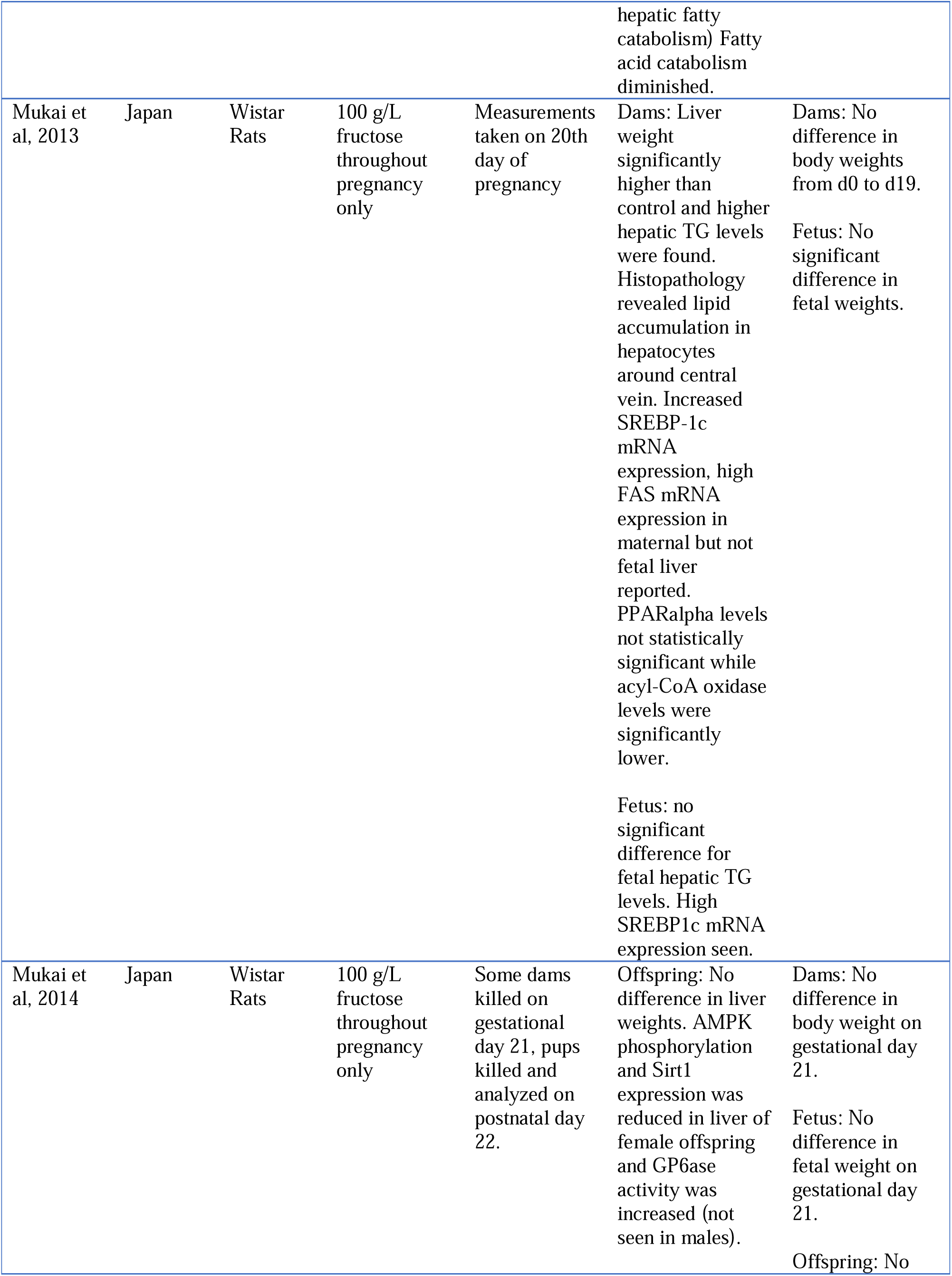
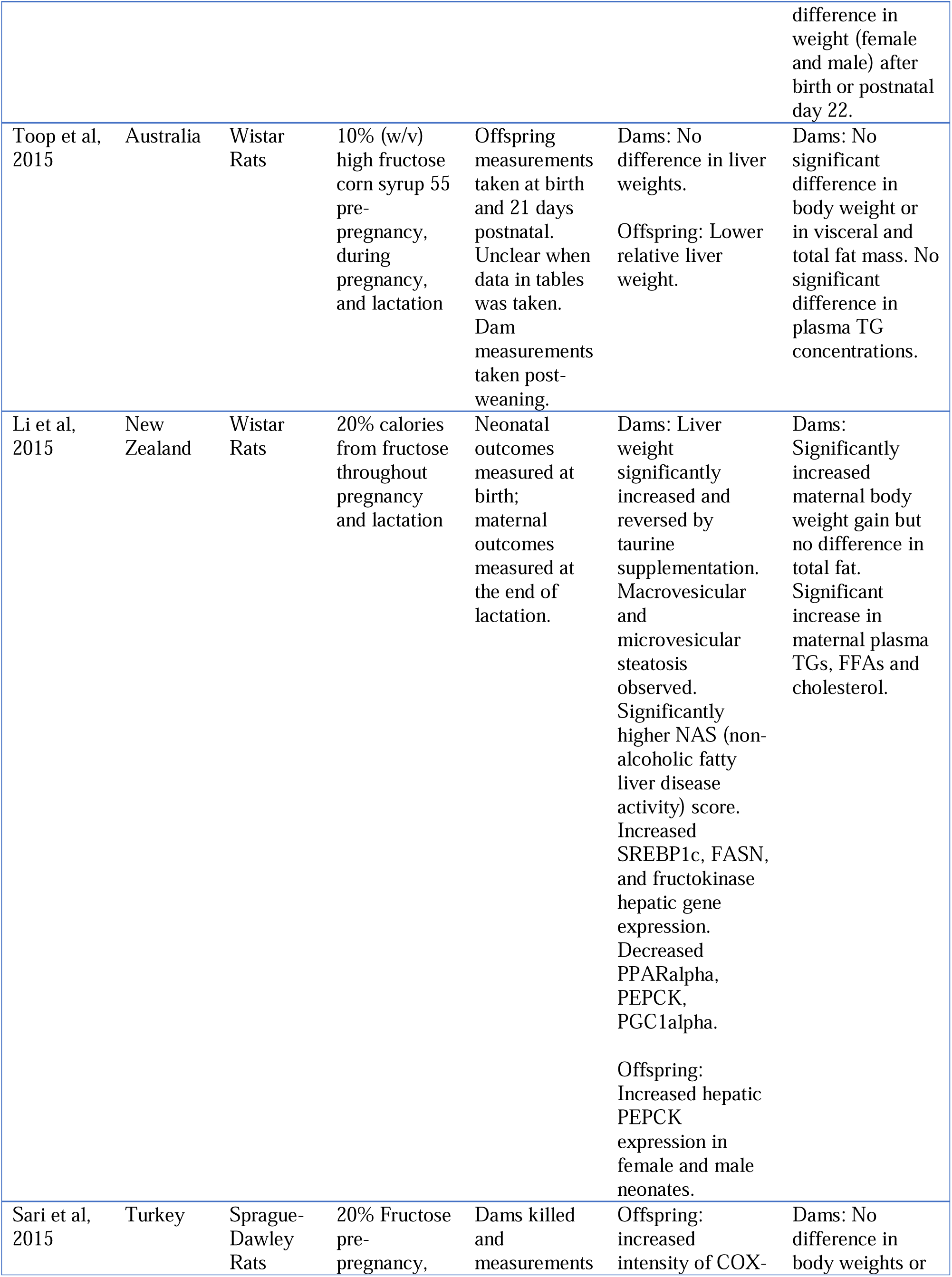
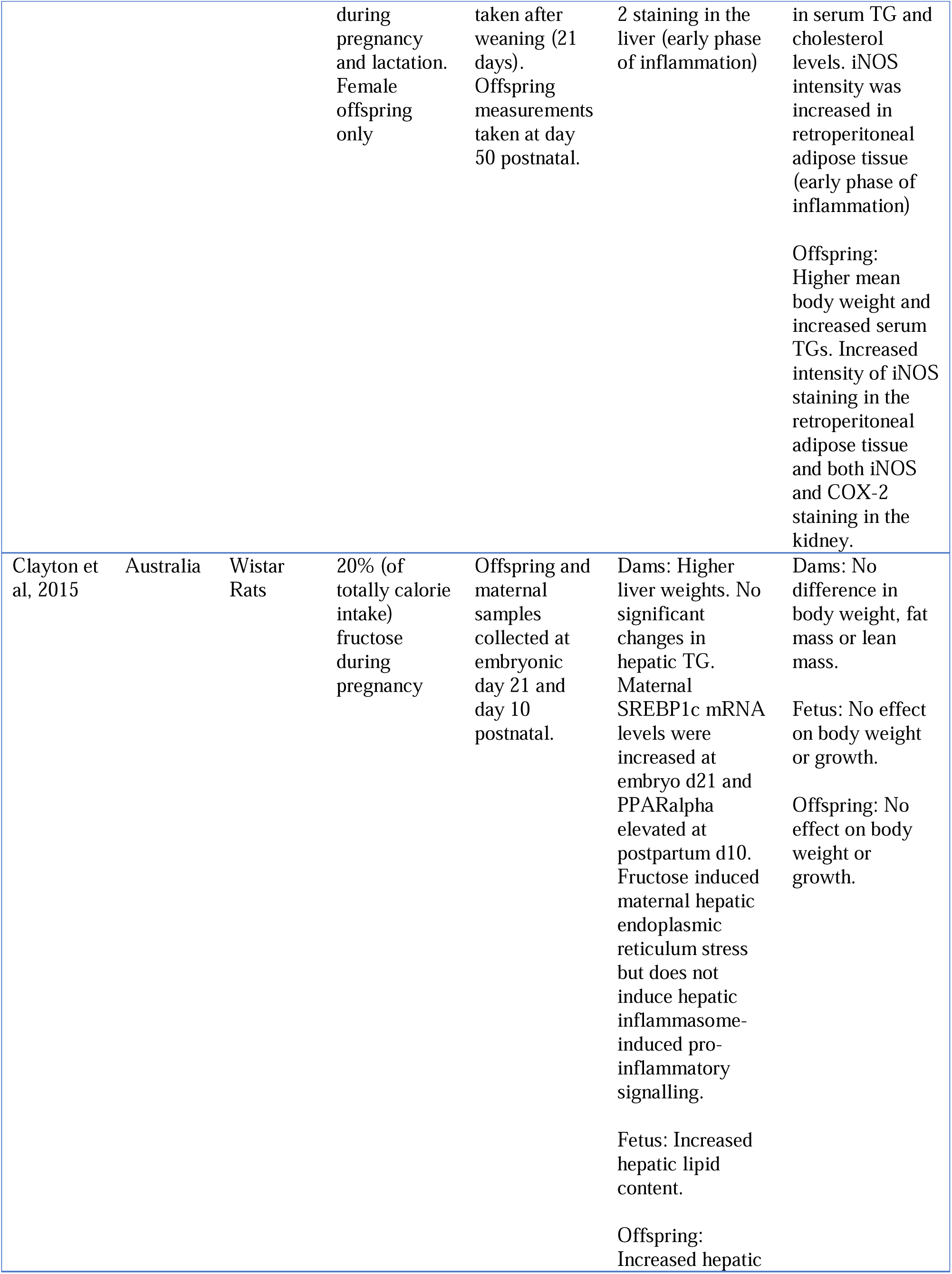
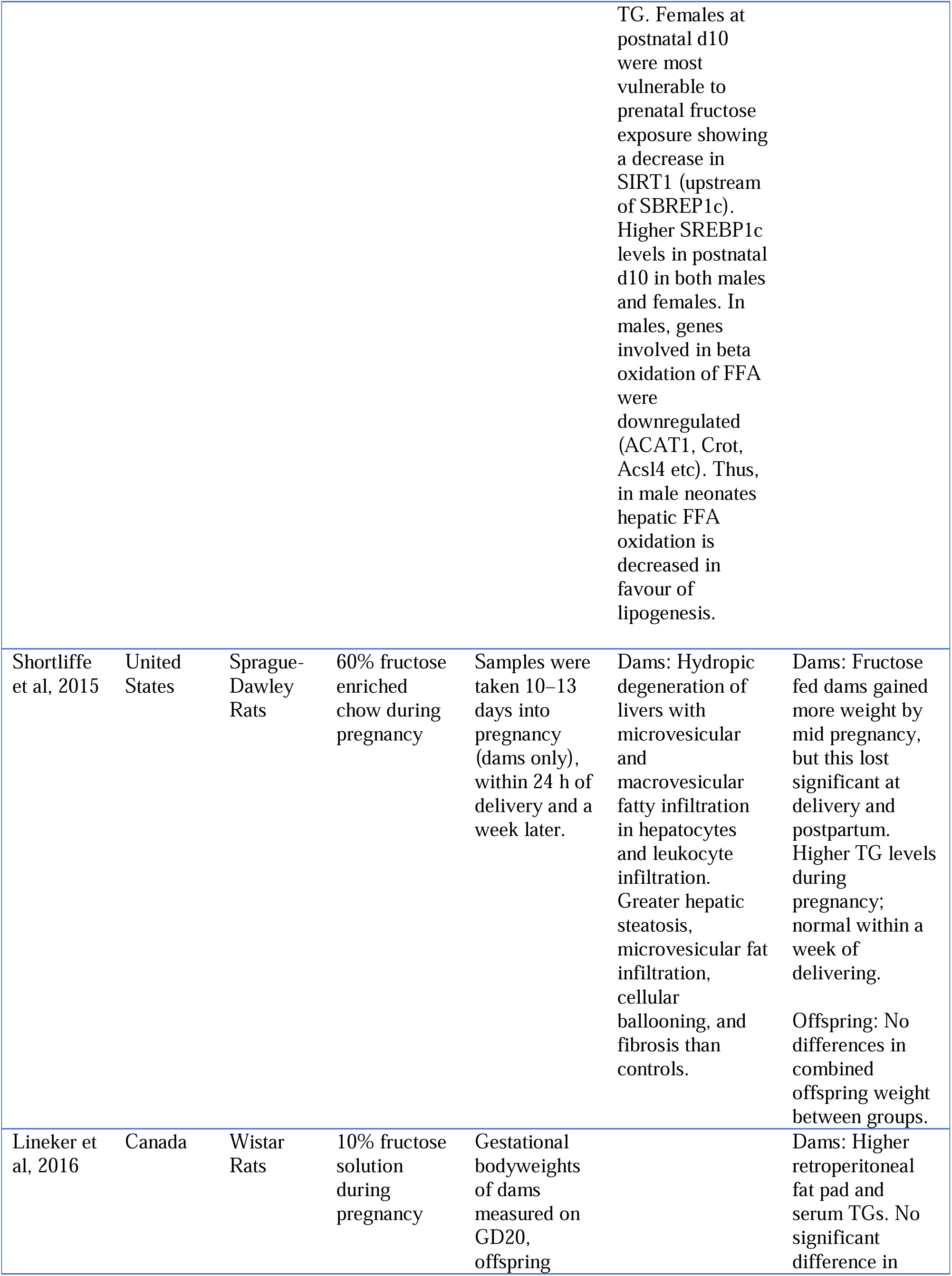
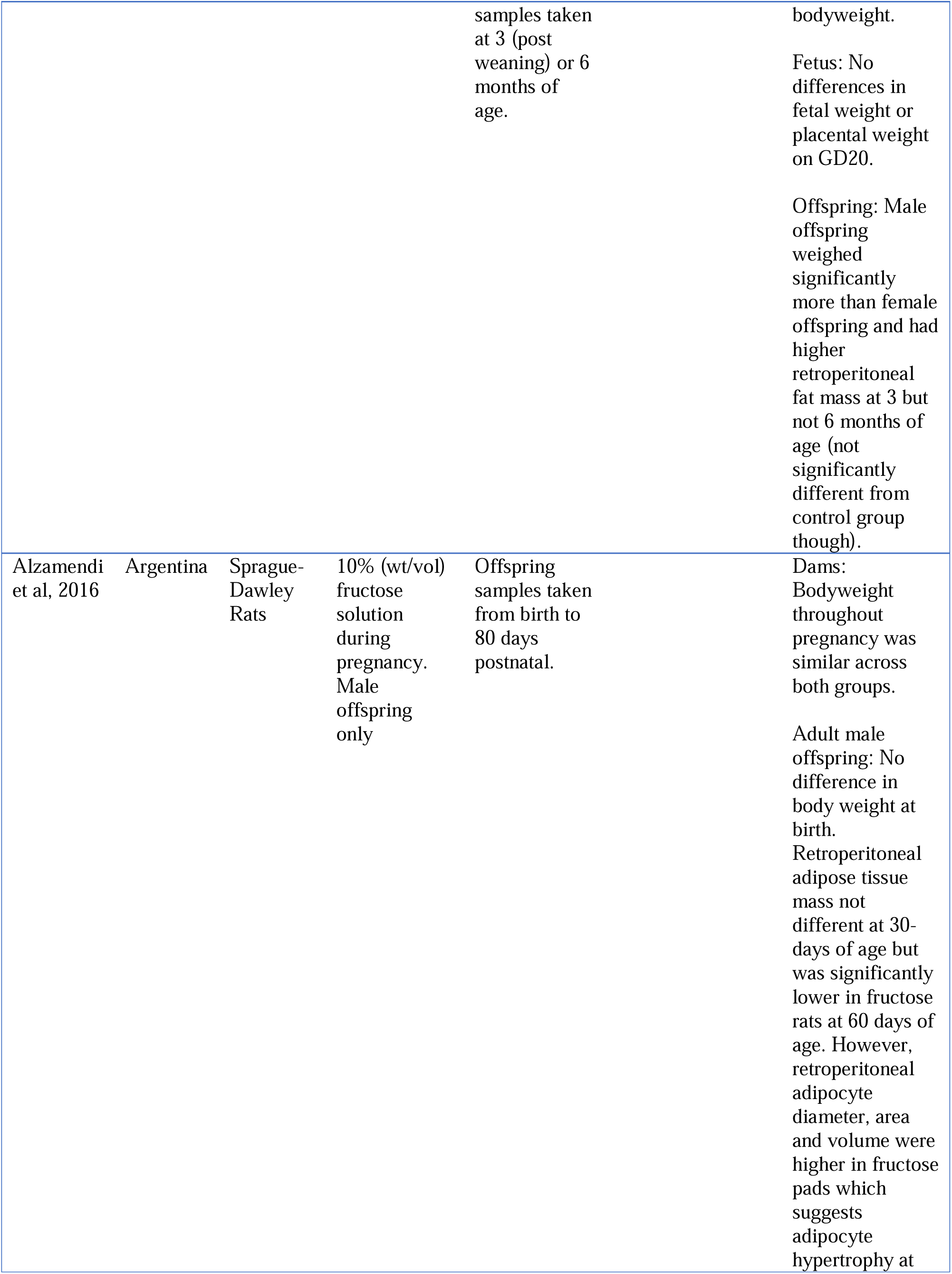
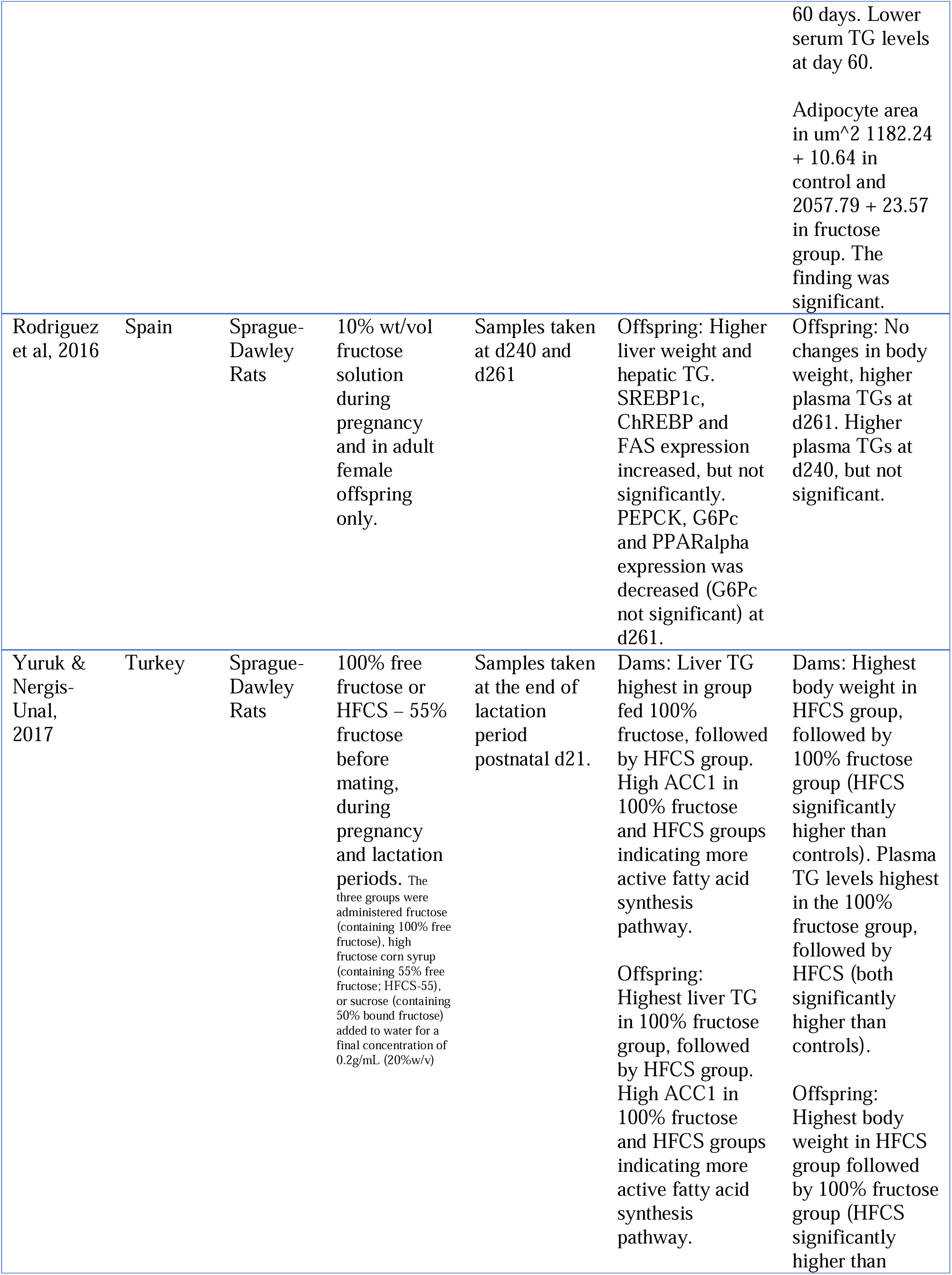
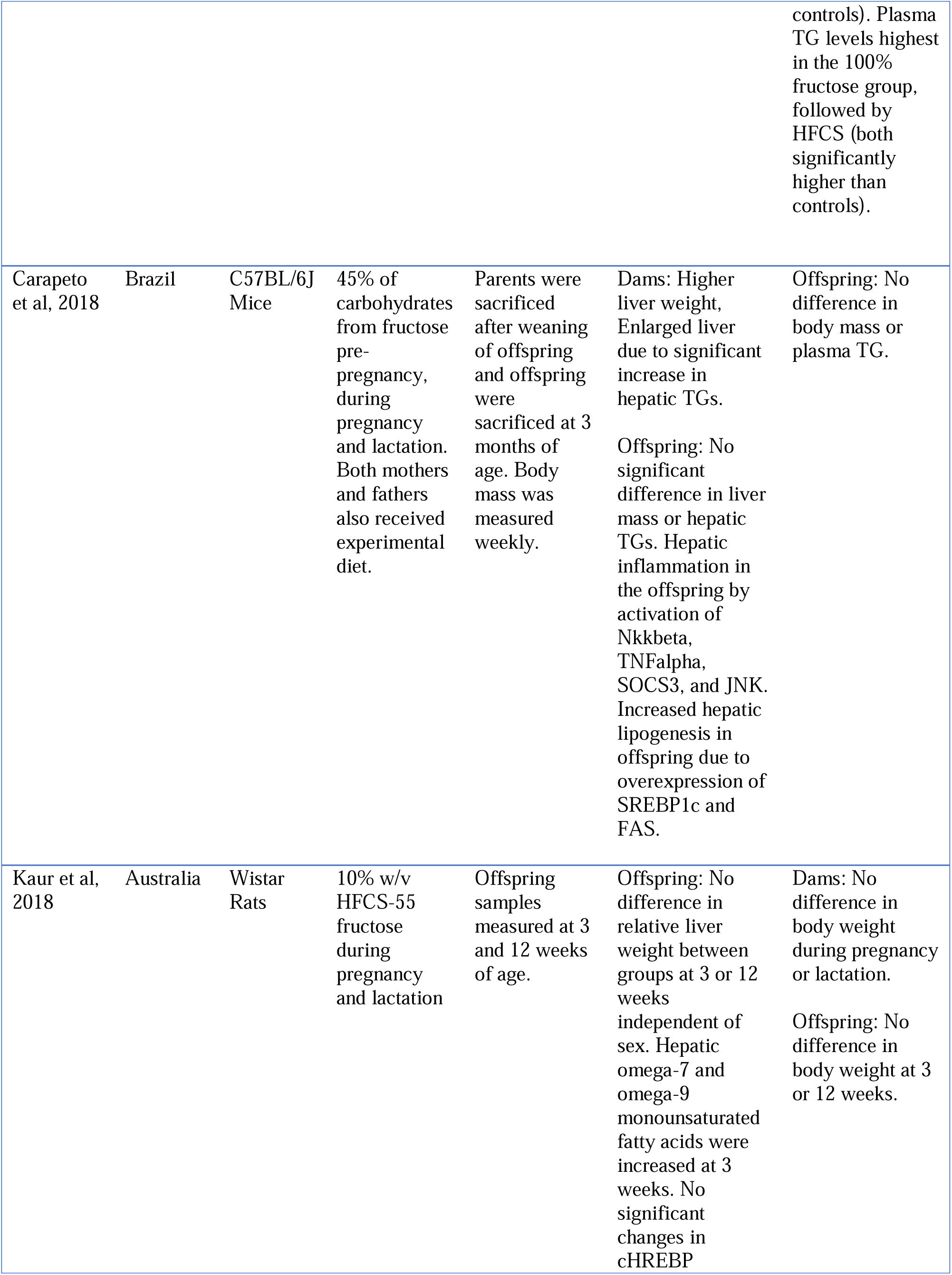
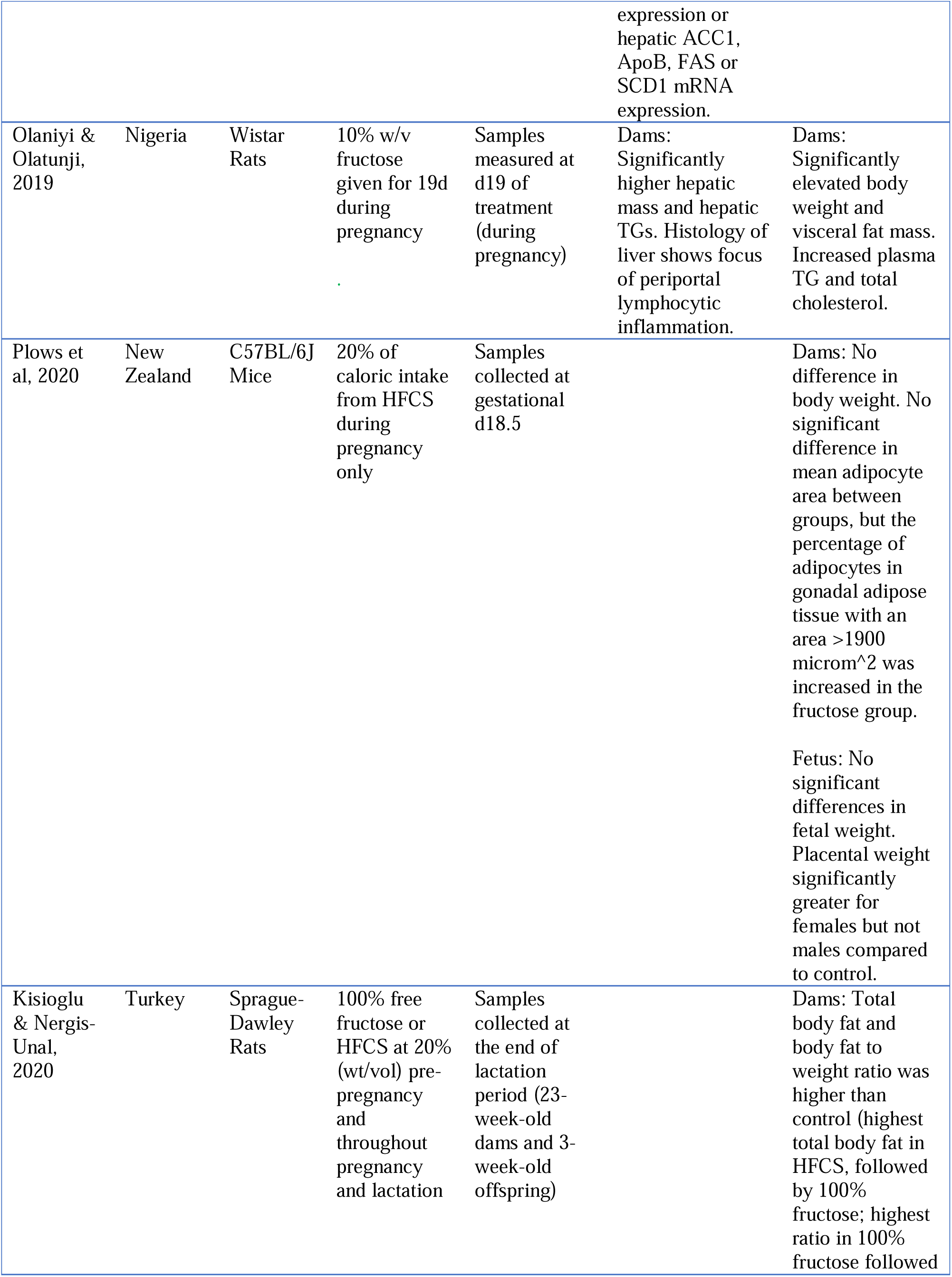
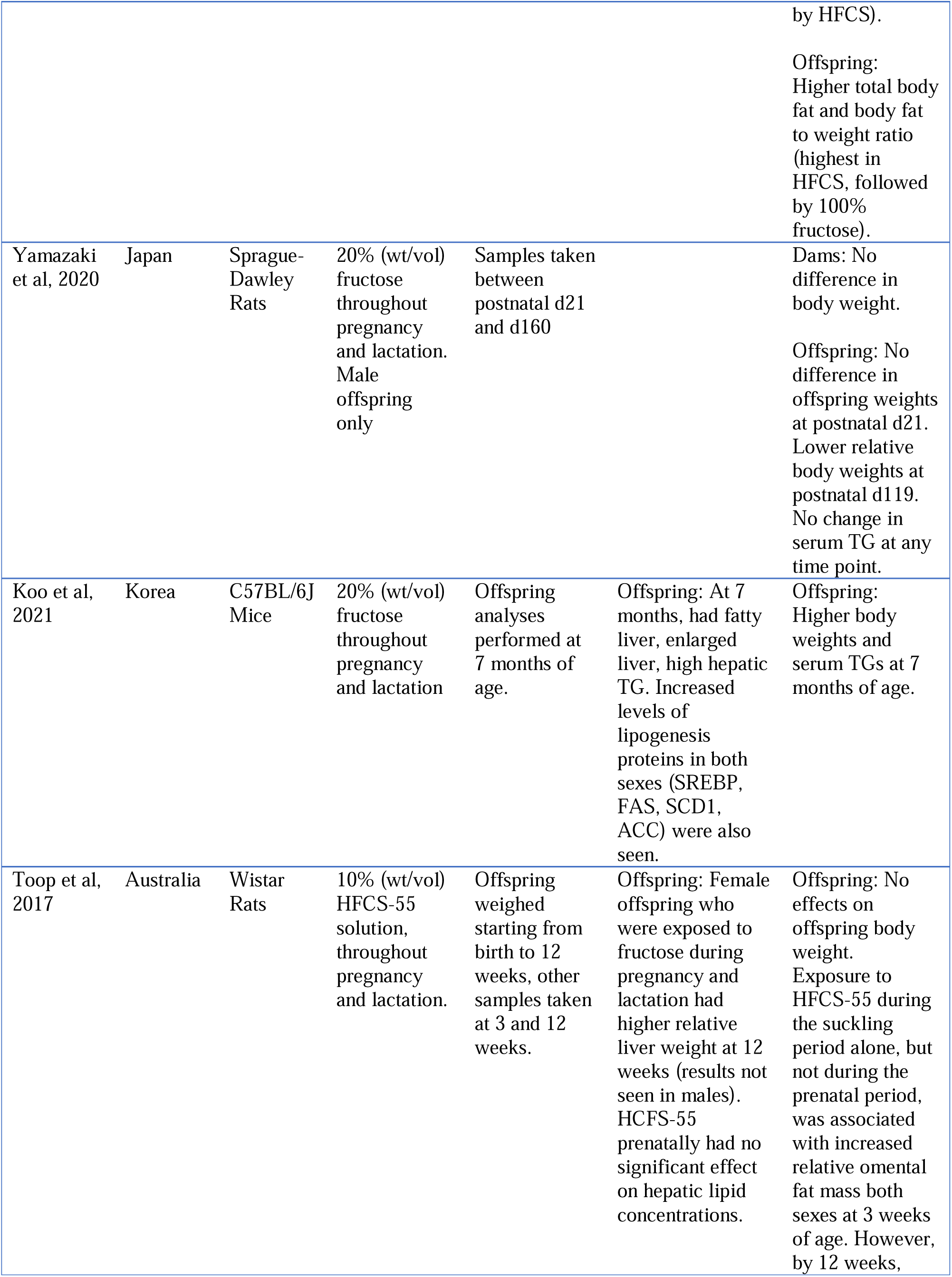
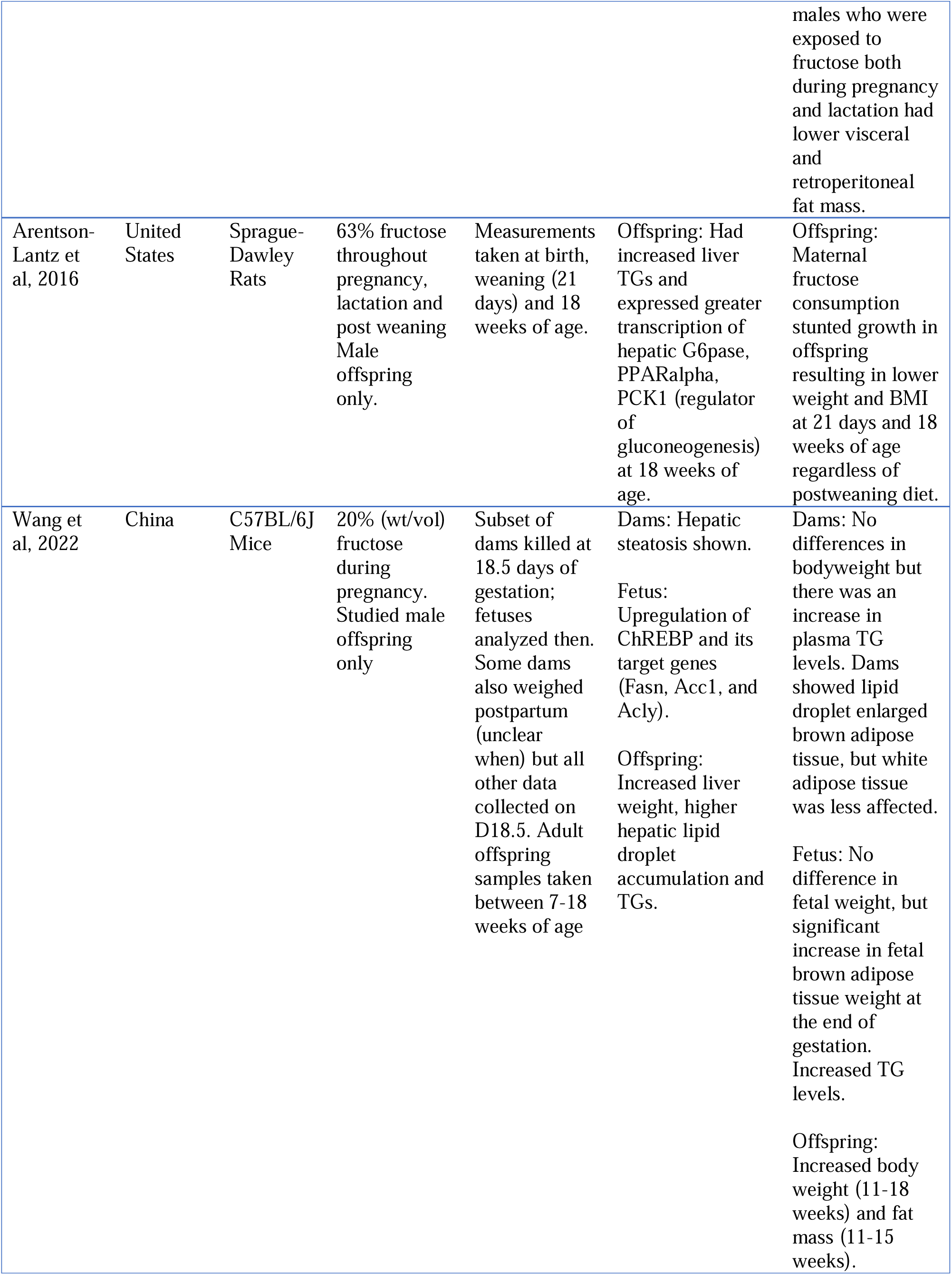
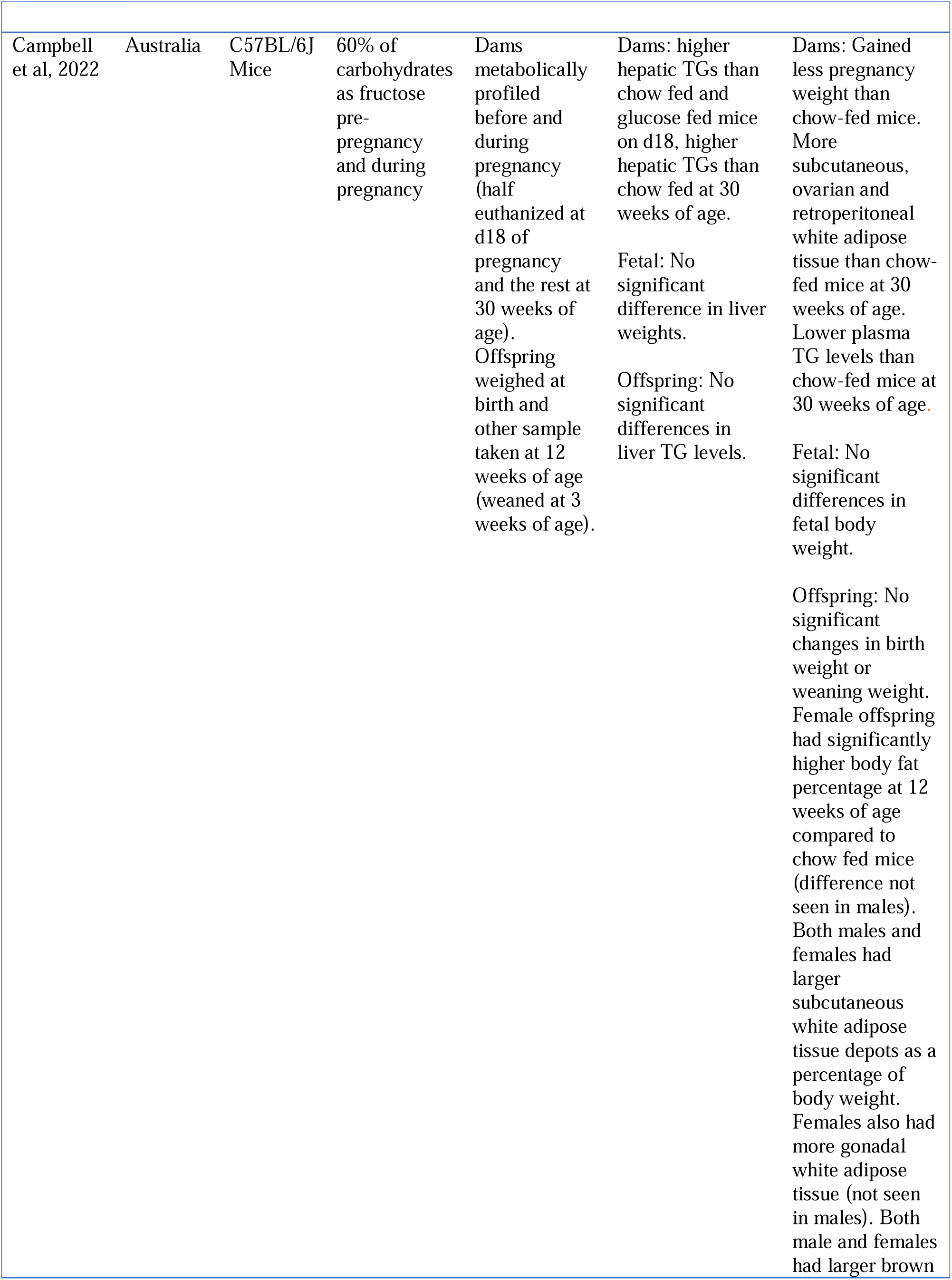
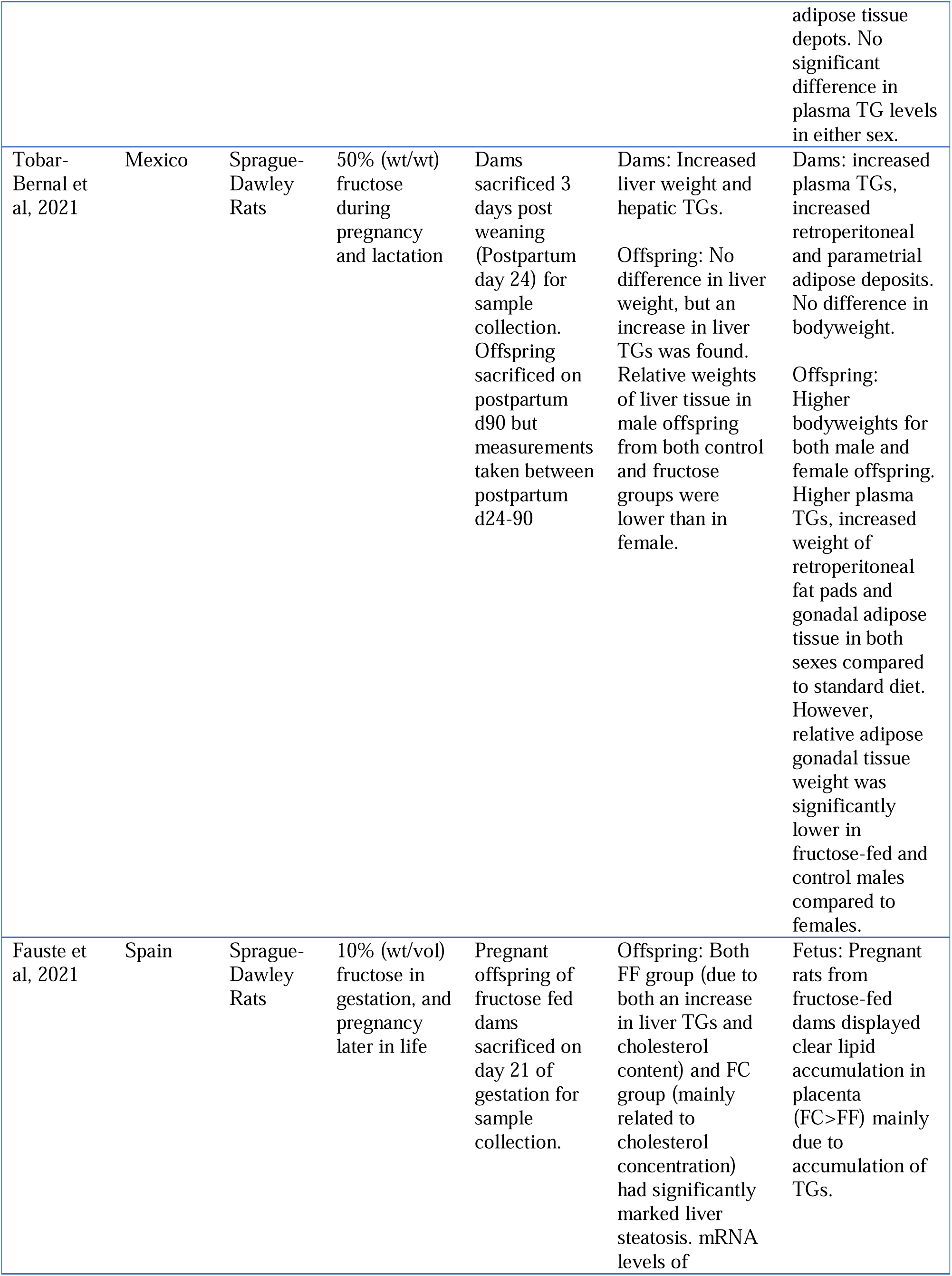
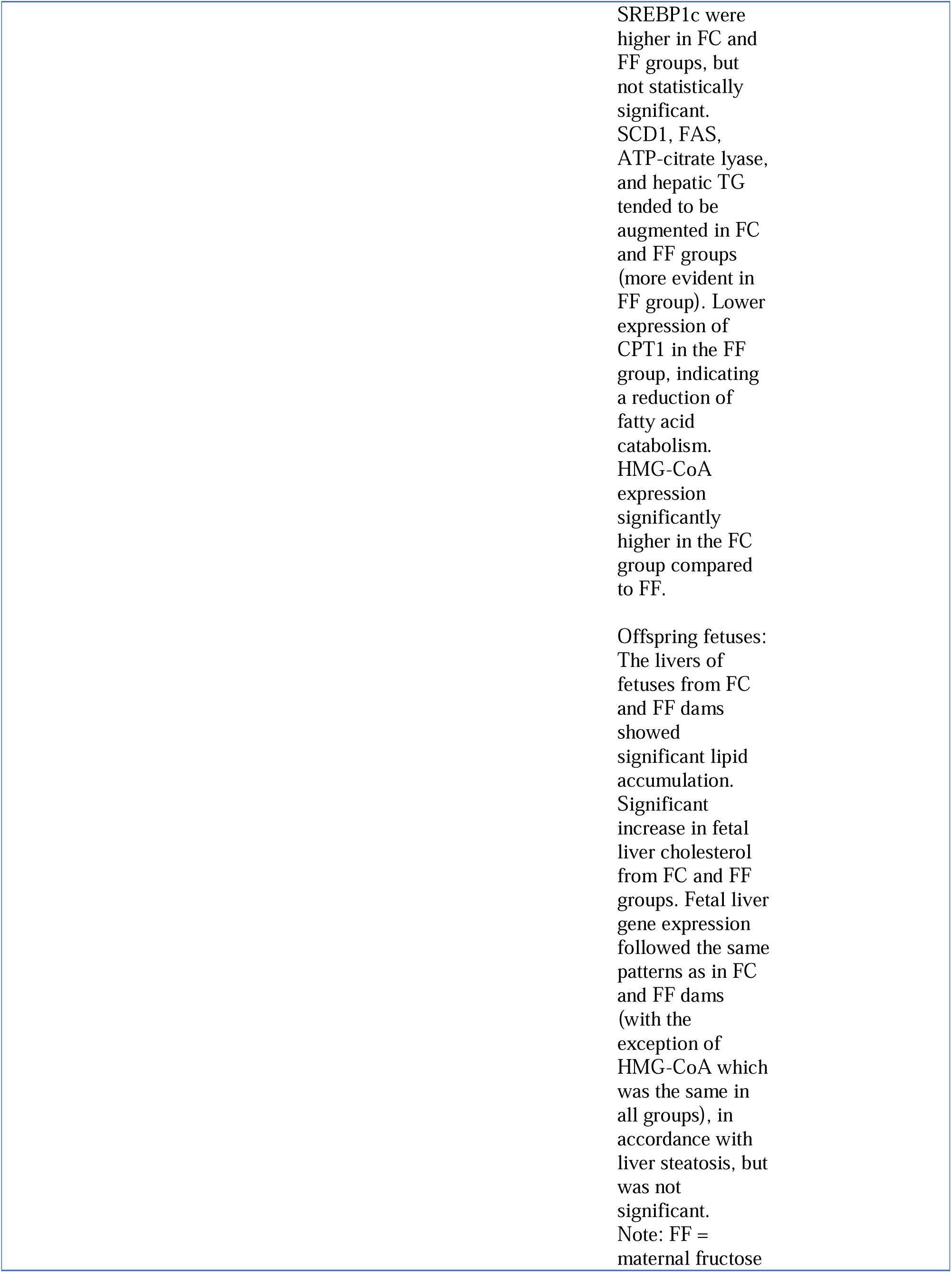
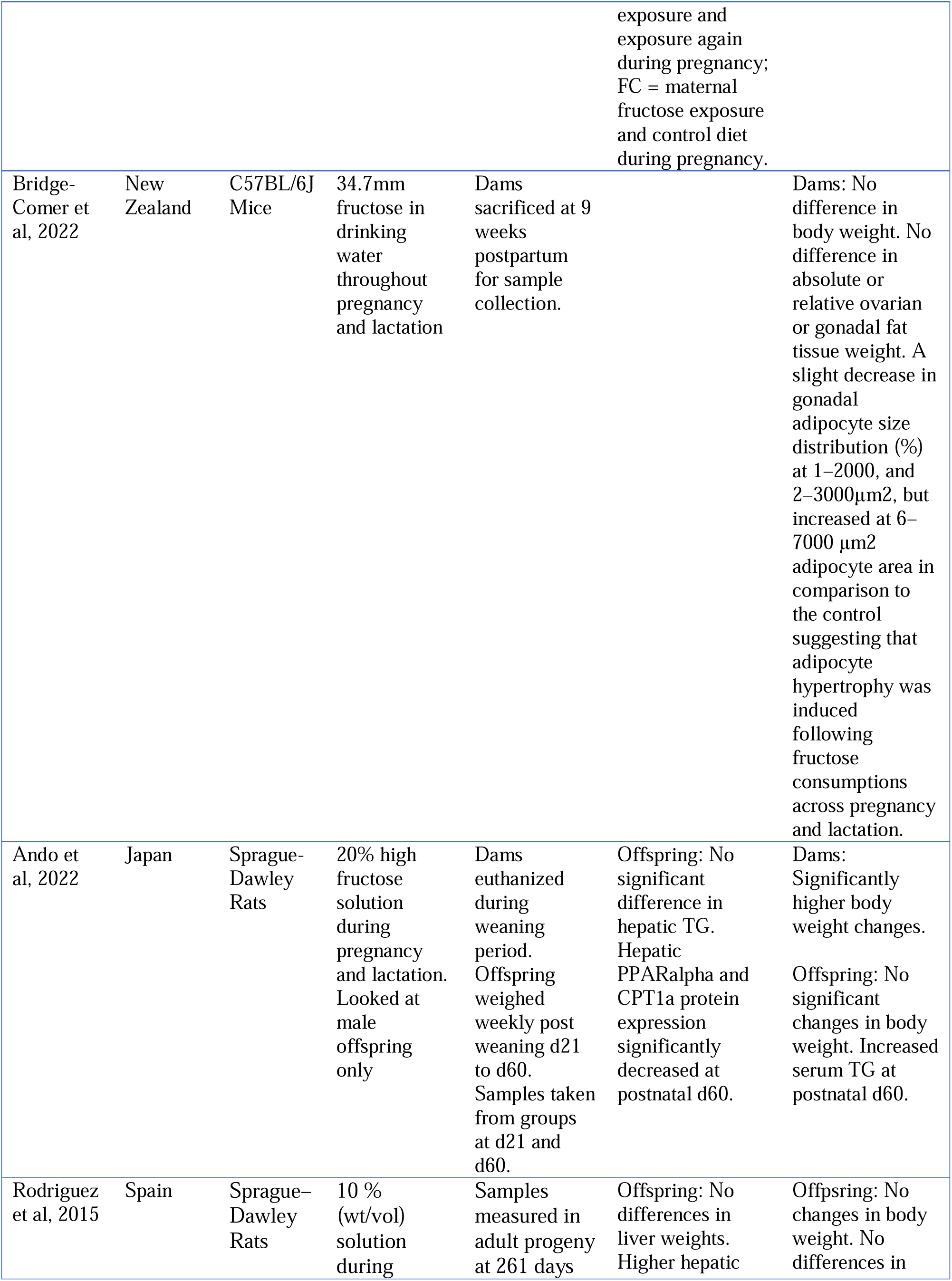
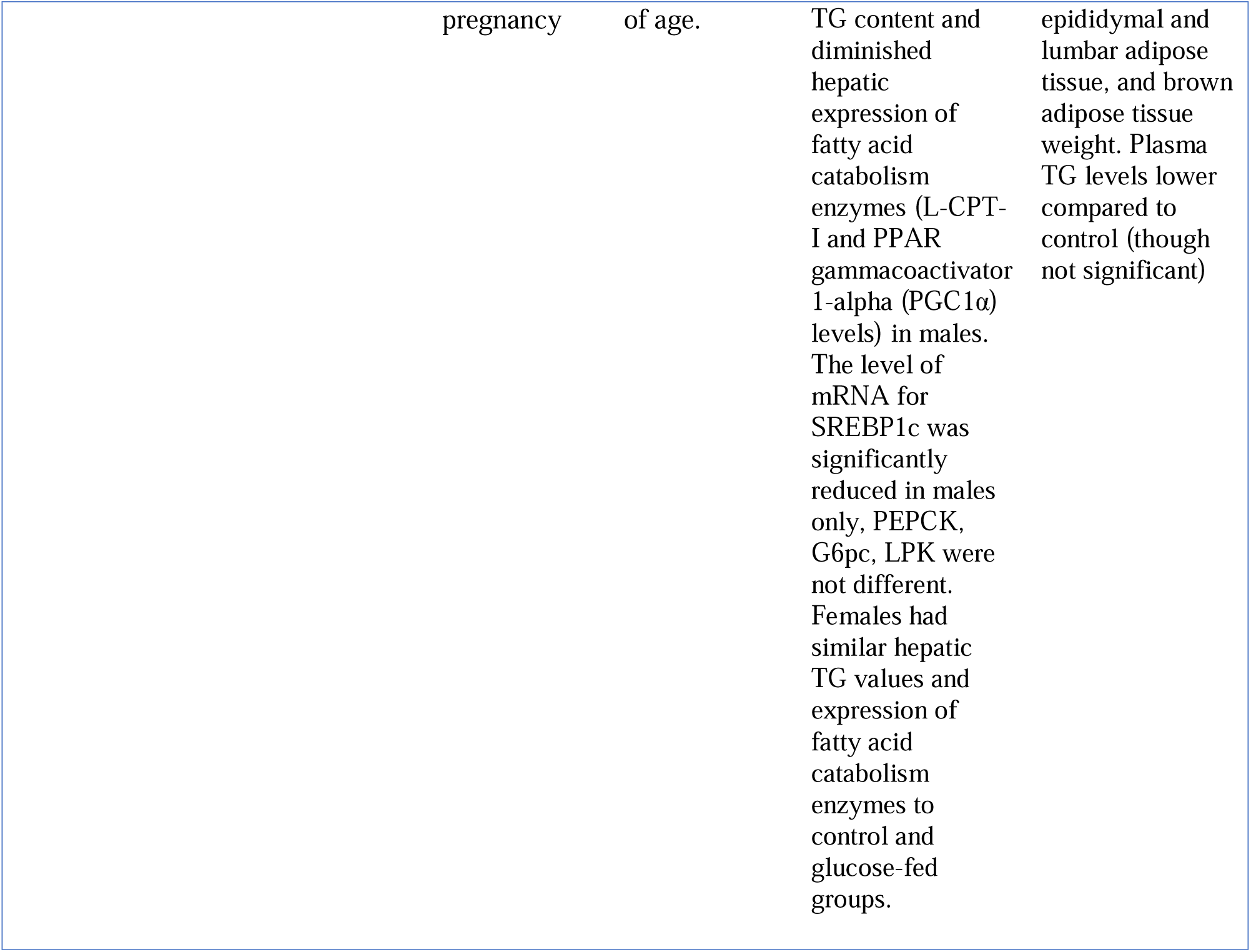
Summary of findings from all included articles (n=33)

### Whole-Body Adiposity

#### Maternal Outcomes

While two studies recorded a decrease in body weight in fructose-fed dams^45, 77^ and others reported an increase in body weight,^61, 63, 73, 75, 78^ most (18 out of 25 studies) reported no difference in body weight in fructose-fed dams compared to control groups.^42, 43, 52, 54-56, 59, 62, 65, 66, 68-70, 72, 74, 79, 80^ 9 of the 12 studies that discussed whole-body adiposity supported our definition of maternal whole-body adiposity in fructose-fed dams (**Table 3**). Specifically, studies reported increases in total body fat mass,^60, 73^ adipocyte hypertrophy,^80^ retroperitoneal,^52, 66, 77^ visceral,^61, 77^ or parametrial hypertrophy,^52^ or mean gonadal adipose tissue area,^79^ lipid droplet enlarged brown adipose tissue^42^ or ovarian adipose tissue mass.^77^ However, four studies reported no difference in various body fat measures including fat mass or total fat percentage,^43, 69^ visceral adipose tissue mass,^69^ mean adipocyte area,^79^ gonadal or ovarian adipose tissue weight^80^ in fructose-fed dams. Of note, it was a common finding for fructose-fed dams to have increased serum^55, 66, 72^ and plasma^42, 52, 54, 56, 61, 73, 78^ triglyceride levels during pregnancy, without changes in body composition.

**Table 3:** Summary of studies supporting maternal whole-body adiposity results (n=12)

| Association with Whole-Body Adiposity | Fructose Exposure (wt:vol or wt:wt) |  |  |  |  |
| --- | --- | --- | --- | --- | --- |
|  |  |  | Other: |  |  |
|  | < 30%, low dose | > 30%, high dose | 20% (of total calorie intake) | 60% (of carbohydrates) | 34.7mm fructose in drinking water |
| **Yes (n=9)** | Rawana et al, 1993; Lineker et al, 2016; Olaniyi & Olatunji, 2019; Kisioglu & Nergis-Unal, 2020; Wang et al, 2022. | Tobar-Bernal et al, 2021. | Plows et al, 2020. | Campbell et al, 2022. | Bridge-Comer et al, 2022. |
| **No (n=3)** | Toop et al, 2015. |  | Clayton et al, 2015; Li et al, 2015. |  |  |

#### Fetal *Outcomes*

None of the eight studies reporting fetal weight found differences between control and maternal fructose-exposed fetuses.^42, 43, 66, 70-72, 77, 79^ Interestingly, one study reported a significant increase in fetal brown adipose tissue weight at the end of gestation, despite no change in overall fetal body weight.^42^ Additionally, this study also reported higher fetal ChREBP mRNA expression and its target genes (FASN, ACC1, and ACLY) in fetal brown adipose tissue.^26^

In a transgenerational study, pregnant rats from fructose-fed dams exhibited significant placental lipid accumulation, primarily due to triglyceride buildup, regardless of whether they were fed fructose or control diets during their own pregnancy.^57^

#### *Offspring* Outcomes

Similar to maternal outcomes, most studies (n=17) reported no significant differences in offspring weight at various time points for fructose-fed dams (Table 2).^43, 45, 53-55, 59, 62, 64-67, 70, 74-77, 81^ Six out of 10 studies found significant increases in offspring measures of adiposity^42, 52, 60, 65, 74, 77^ (**Table 4**). These studies found increases in total body fat,^60, 77^ brown adipose tissue (in both sexes),^77^ visceral adipose tissue,^74^ fat mass,^42^ subcutaneous fat tissue,^77^ retroperitoneal,^52^ and gonadal adipose tissue.^52, 77.^ One study reported reduced retroperitoneal adipose tissue mass but increased adipocyte hypertrophy in male offspring at 60 days postnatal (Table 2).^65^ Three studies identified no significant difference in fat content compared to control groups,^66, 67, 73^ One study found that offspring exposure to high-fructose corn syrup only during the suckling period, not the prenatal period, increased relative visceral adipose tissue mass at three weeks.^64^

**Table 4:** Summary of studies supporting offspring whole-body adiposity results (n=10)

| Association with Whole-Body Adiposity | Fructose Exposure (wt:vol or wt:wt) |  |  |  |  |
| --- | --- | --- | --- | --- | --- |
|  | Other: |  |  |  |  |
|  | < 30%, low dose | > 30%, high dose | 20% (of total calorie intake) | 45% (of carbohydrates) | 60% (of carbohydrates) |
| **Yes (n=6)** | Saad et al, 2016; Alzamendi et al, 2016; Kisioglu & Nergis-Unal, 2020; Wang et al, 2022. | Tobar-Bernal et al, 2021. |  |  | Campbell et al, 2022. |
| **No (n=4)** | Rawana et al, 1993; Lineker et al, 2016; Toop et al, 2017; Rodriguez et al, 2015. |  |  |  |  |

Like the observations of TG levels in maternal circulation, results in offspring varied regarding serum triglyceride levels. Seven studies reported higher plasma or serum triglyceride levels in the offspring of fructose-fed dams,^52, 55, 58, 68, 73, 75, 76^ compared to the four that reported lower levels^65^ or no effect.^59, 77, 81^

### Hepatic Adiposity

#### Maternal *Outcome*

In ten studies, fructose-fed dams were found to have higher liver weight.^43, 45, 52, 55, 61, 71, 73, 76, 78, 81^ Only one study reported no change in liver weights^69^ and one reported lower liver weights.^56^ In the majority of these reports, low-dose fructose consumption (<30% wt:vol or wt:wt) was used (**Table 5**).

**Table 5:**
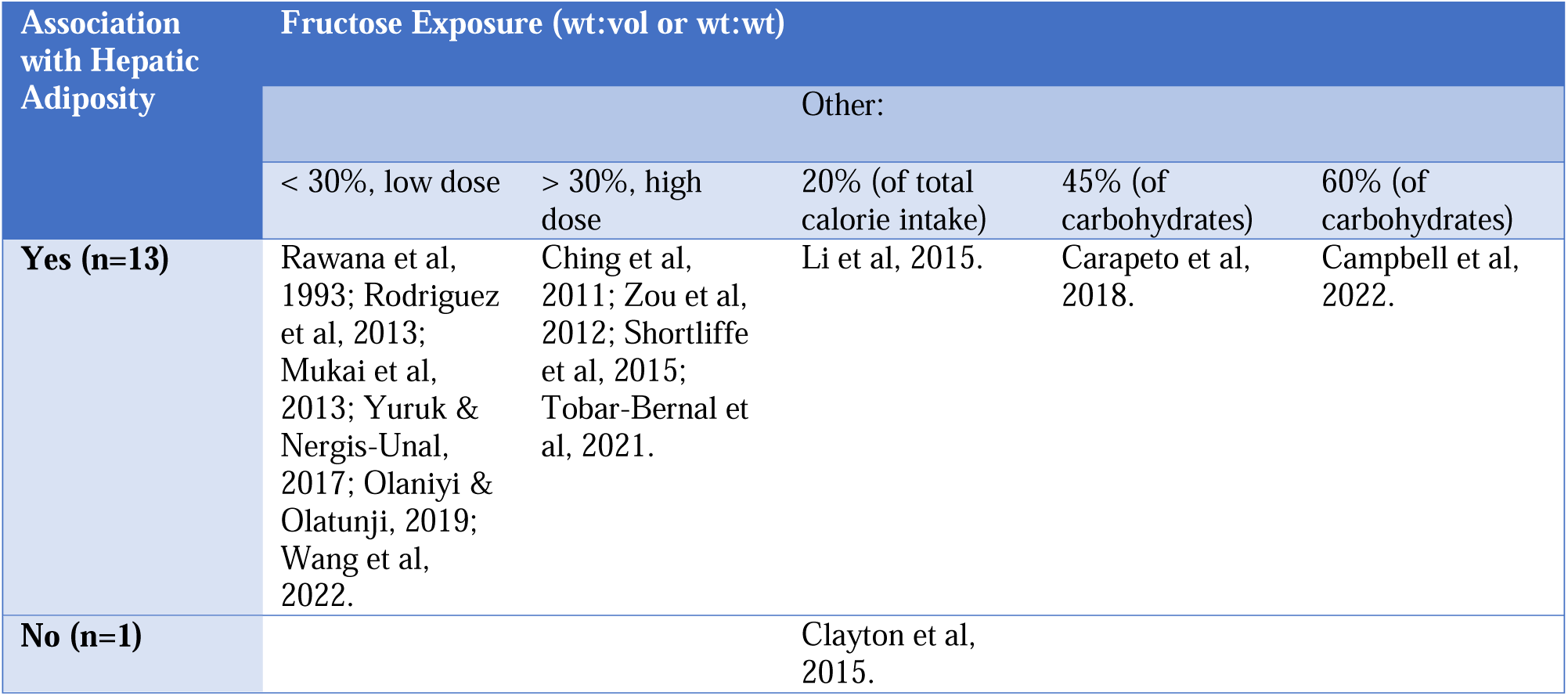
Summary of studies supporting maternal hepatic adiposity results (n=14)

From 14 studies that did investigate hepatic lipid accumulation after prenatal fructose exposure, 13 studies supported the association between maternal fructose consumption and maternal hepatic adiposity (Table 5). Specifically, studies reported higher levels of hepatic triglycerides,^45, 52, 55, 61, 63, 71, 72, 77, 81^ hepatic lipid concentrations,^73^ and hepatic steatosis^42, 54, 72, 78^ in fructose-fed dams. In papers that reported histological findings of the liver periportal lymphocytic inflammation^61^ and lipid droplets in periportal hepatocytes^45, 54, 71^ were observed.

Elevated mRNA levels of associated with key lipid synthesis providing potential mechanisms, associated with increased maternal fructose consumption, leading to elevated maternal hepatic adiposity. This was associated with elevated gene expression of enzymes involved in fatty acid synthesis (e.g. ChREBP, FAS, acetyl-coA carboxylase)^71, 72, 78^ as well as SREBP1c.^43, 71, 78^ Additionally, PEPCK gene expression, a rate-limiting enzyme in gluconeogenesis, was elevated in the livers of fructose-fed dams.^45, 56, 73^ Of note, one study reported no significant changes in maternal hepatic triglyceride levels, however, both maternal and offspring SREBP1c mRNA levels were increased, without increased hepatic lipid accumulation.^43^

#### *Fetal* Outcomes

Six studies reported effects of prenatal fructose exposure on the fetal hepatic outcomes (Table 2).^42, 43, 57, 71, 72, 77^ Of these studies five, did not report fetal liver weight,^26, 43, 57, 71, 72^ one study reported no significant difference in liver weight.^77^ Three studies reported fetal hepatic steatosis^43, 57, 72^ while one study reported no significant differences in hepatic triglyceride levels, although expression of hepatic lipogenic proteins (eg. SREBP1c and SCD1) were elevated.^71^ One study reported an upregulation of ChREBP and its target genes (*e.g*., FASN, ACC1, and ACLY).^26^

#### Offspring *Outcomes*

A decrease,^69^ or no change^52, 55, 62, 67, 70, 81^ in liver weight was reported in the offspring of fructose-fed mothers in seven studies. Four studies documented an increased liver weight^42, 64, 76^ or enlarged liver^58^ in the offspring. Interestingly, hepatic adipose content varied depending on the age of the offspring, with one study reporting increased adipose content at 3 weeks of age, but no significant changes at 12 weeks.^64^ Additional information on the age of data collection and results obtained at these times can be found in Figure 2 and Table 2.

14 out of 17 studies that investigated hepatic fat accumulation in the offspring of fructose-fed dams found hepatic steatosis in the offspring (Table 6).^42, 43, 52, 53, 55, 57, 58, 62, 63, 67, 68, 74, 76, 81^ Hepatic adipose accumulation^42, 43, 52, 53, 55, 57, 58, 62, 63, 67, 68, 74, 76, 81^ was commonly found amongst offspring of fructose-fed dams, in association with increased expression of lipogenic proteins such as SREBP1c and FAS^43, 57, 71, 76, 81^. With four studies reporting no difference in offspring hepatic triglyceride levels.^64, 75, 77, 81^ Interestingly, Fauste *et al* (2021) reported an intergenerational effect where pregnant offspring of fructose-fed dams had significantly marked hepatic steatosis in both fructose diet and control diets during pregnancy.^57^

**Table 6:** Summary of studies supporting offspring hepatic adiposity results (n=17)

| Association with Hepatic Adiposity | Fructose Exposure (wt:vol or wt:wt) |  |  |  |  |
| --- | --- | --- | --- | --- | --- |
|  | Other: |  |  |  |  |
|  | < 30%, low dose | > 30%, high dose | 20% (of caloric intake) | 45% (of carbohydrates) | 60% (of carbohydrates) |
| **Yes (n=14)** | Saad et al, 2016; Rodriguez et al, 2016; Yuruk & Nergis-Unal, 2017; Koo et al, 2020; Wang et al, 2022; Fauste et al, 2021; Rodriguez et al, 2015; Sari et al, 2015; Kaur et al, 2018. | Ching et al, 2011; Arentson-Lantz et al, 2016; Tobar-Bernal et al, 2021. | Clayton et al, 2015. | Carapeto et al, 2018. |  |
| **No (n=3)** | Toop et al, 2017; Ando et al, 2022. |  |  |  | Campbell et al, 2022. |

#### *Sex* Differences

Several studies report sex differences in hepatic adipose content, liver weight, and offspring weight. Female offspring of dams exposed to 10% wt/vol fructose during pregnancy had higher liver fat infiltrates, greater offspring weight at one year, and increased relative liver weights compared to male offspring. ^64,74^ Similarly, one study found a significant reduction in male, but not female fetal body weight with an associated increase in placental weight in females.^79^ Female offspring of fructose-fed mice had significantly higher body fat percentage at 12 weeks of age compared to chow-fed mice and had more visceral adipose tissue at one year of age when compared to their male counterparts.^74, 77^ In contrast, one study found that male offspring weighed significantly more than female offspring and had higher retroperitoneal fat mass at 3 months of age.^66^ The higher liver weight found in female offspring of fructose-fed dams in the paper by Rodriguez *et al* (2016) was associated with increased mRNA expression of SREBP1c, ChREBP, and FAS.^76^ In another study, hepatic mRNA level of SREBP1c in male offspring was significantly reduced.^67^

Sex differences have also observed in interventional studies, aimed at mitigating the effects of excessive fructose feeding. Li *et al*. (2015) found that maternal taurine supplementation attenuated maternal fructose-induced hepatic tumour necrosis factor receptor 1in male offspring, but not female offspring.^78^ Taurine supplementation also did not appear to effect the elevated hepatic PEPCK mRNA expression in female offspring, although it did normalize it in male offspring.^78^ Furthermore, the expression of SIRT1, an enzyme that inhibits lipogenesis, was significantly reduced in female offspring but not in males.^43, 70^

## DISCUSSION

The current scoping review has revealed that fructose consumption during pregnancy in rodent models is associated with significant increases in whole-body adiposity in dams and offspring. Moreover, maternal fructose consumption was associated with hepatic steatosis in dams, fetuses, and offspring. Several studies identified sex-specific findings where female offspring were found to be more susceptible to prenatal fructose exposure than males, with greater hepatic adiposity and whole-body adiposity. Interestingly, consumption of a high fat/high fructose diet is commonly associated with a reduced or unchanged body weight in guinea pigs, mice, and rats.^82-86^ Although *ad libitum* access to fructose-containing diets do not necessarily increase body weights, it consistently increases fat mass in rodents.^87^ These observations are consistent with our current findings, as most of our studies found no difference in body weight in dams and offspring, however several papers reported adipocyte hypertrophy and adipose tissue mass increases. Our scoping review also reinforced the association between prenatal fructose exposure and hepatic steatosis in rodent dams, fetuses, and offspring. Key regulatory proteins identified in association with this increased hepatic steatosis included ones underlying mechanisms associated with lipid synthesis including ChREBP,^42, 72, 76^ FAS,^57, 58, 71, 72, 76, 81^ and SREBP1c^58, 71, 76, 81, 88^ and their target genes. Thus, the upregulation of networks involved in hepatic *de novo* lipogenesis may likely play a role in the progression of NAFLD in the mother and offspring.

Another potential mechanism in the adverse effects of fructose may be related to the metabolic effects of uric acid, a by-product of fructose metabolism.^89^ Hyperuricemia is a key component of metabolic syndrome.^4, 5^ Interestingly, GLUT9 knockout in enterocytes in mice induces hyperuricemia and elevated hepatic triglycerides and free fatty acids.^10^ High-fructose feeding in guinea pigs prior to and during pregnancy, resulted in offspring with significantly altered serum free fatty acids, and increased levels of uric acid and triglycerides, all of which had developed before weaning.^90^ Additionally, high-fructose (60% fructose) consumption in mice has been demonstrated to increases placental fructose transport *via* GLUT9, and subsequent *de novo* uric acid production by activating the activities of the enzymes AMP deaminase and xanthine oxidase, which resulted in increased lipids and oxidative stress in the placenta.^38, 91^ Treatment with allopurinol, a xanthine oxidase inhibitor, attenuated such effects.^10, 38^ Given these studies highlight an unfavourable impact by excess fructose on various organ systems in the perinatal period, further research is required to elucidate the relationship between fructose metabolism and uric acid production during the perinatal period and later life adipogenesis.

Several studies highlighted that fetal sex may be an important consideration in understanding fructose exposure effects. Sex-specific fetal programming may indicate that the placenta functions in a sex-specific manner.^92, 93^ Interestingly, mice placental weights were found to be significantly greater in females when compared to males, most likely from lipid accumulation.^79^ In contrast, one study found that a high maternal fructose (20% fructose) in rats was associated with decreased placental weights in female fetuses when compared to males.^94^ Female mice offspring tended to develop obesity, had increased adiposity and hepatic lipid deposition following maternal exposure to high fructose diet, whereas male offspring were hypertensive and developed insulin resistance.^74^ It is possible that sex hormones may play a role, however further investigation is warranted.^95^

The current review has also highlighted that there is a lack of data concerning the potential transgenerational impacts of excessive maternal fructose consumption. Rodriguez *et al* (2016) found that low maternal fructose intake (10% wt/vol) in pregnant rats provoked long-term impacts such as impaired insulin signal transduction, hyperinsulinemia, and hypoadiponectinemia in adult male, but not female progeny.^95^ One transgenerational study by Fauste *et al* (2021) found that fructose intake during pregnancy contributes to fetal programming, such that their progeny in future pregnancies develop the same programmed phenotype regardless of fructose exposure.^57^ These reports demonstrate that fructose exposure pre-pregnancy and during pregnancy may have lasting deleterious effects on metabolic health on future generations.

It is challenging to translate the results of animal studies to humans as the timing of development of major organ systems, such as adipose depots and liver, impacted *in utero* differs in humans which are a precocial species, compared to most rodent species which are altricial, where the young are underdeveloped at the time of birth. Second, there is great heterogeneity with respect to the dose of fructose consumed during pregnancy, as well as the timing to which fructose was administered. Fructose 10% wt/vol consumed by rodents over time, mimics how fructose is predominantly ingested in human populations, i.e., fructose-sweetened soft drinks, while, high levels of fructose (30-60% wt/vol) could be deemed supraphysiological, and do not reflect the average human intake.^49, 50, 96^ Furthermore, while some studies reported on liver weight, they did not record any other measures of hepatic accumulation, and as such were not included in the analysis of studies that supported hepatic adiposity.^56, 69^ Lastly, the studies included, investigated fructose independently as opposed to a Western diet, which can limit interpretation of the specific effects of fructose.

## Conclusion

This scoping review has highlighted that fructose consumption prior to and during pregnancy is associated with maternal and offspring whole-body adiposity, hepatic fat accumulation, sex-specific effects and potential transgenerational effects in rodent models. We highlight the need for human based studies and to improve translation, standardised doses and periods of exposure in animal model systems. The work also highlights potential avenues for further research into the sex-specific effects of maternal fructose consumption on the placenta and offspring, and the mechanisms of developmental programming of offspring phenotypes. Given the lack of human studies on this topic, exploring the impact of prenatal fructose consumption in humans on both fetal and maternal health will be crucial to understand the translation of studies using rodent models. In conclusion, investigating the mechanisms of developmental programming induced by maternal fructose exposure is essential for understanding the long-term health implications of dietary habits during pregnancy. Identifying critical windows of susceptibility and molecular pathways involved in the vertical transmission of these effects can inform public health guidelines ultimately reducing the risk of chronic diseases such as NAFLD, obesity, diabetes, and cardiovascular disorders in the offspring and future generations.

## Acknowledgements/Contribution Statement

GZ and TRHR conceptualized and designed the study. MS performed the initial and updated search strategy. GZ, TRHR, SC, CG, and SG conducted the screening process. GZ and SC performed data collection and TRHR verified it. GZ and SC summarized the data. GZ drafted the first version of the manuscript. All authors contributed critically to interpretation of the data and manuscript revisions. All authors had full access to the data in the study and take responsibility for the integrity of the data and accuracy of the analysis. The corresponding author attests that all listed authors meet authorship criteria.

