## Appendix A for "Fructose consumption in pregnancy and associations with maternal and offspring hepatic and whole-body adiposity: a scoping review"

**Fructose & Pregnancy Searches**

**Medline (OVID)**

Search conducted on November 17^th^, 2021:

1463 TOTAL results

Updated search conducted on August 23^rd^, 2022:

113 results from search/58 duplicates removed by Covidence/ 55 TOTAL new Medline results added.

**Embase (OVID)**

Search conducted on November 22nd, 2021:

1679 results from search/924 duplicates removed by Covidence/755 TOTAL Embase results.

Updated search conducted on August 23^rd^, 2022:

92 results from search/79 duplicates removed by Covidence/ 13 TOTAL new Embase results added.

**Cochrane Library**

Search conducted on November 22^nd^, 2021:

84 results from search/ 36 duplicates removed by Covidence/ 48 TOTAL results from Cochrane.

Ovid MEDLINE

1	fructose.tw. or High Fructose Corn Syrup/ or Fructose/

2	"High Fructose Corn Syrup".tw.

3	pregnancy/ or maternal-fetal exchange/ or placentation/

4	(pregnancy or "maternal-fetus exchange" or placenta*).tw.

5	Placenta/

6	Fetus/ or fetus.tw.

7	neonat*.tw. or Infant, Newborn/

8	offspring.tw.

9	Maternal Exposure/ or maternal.tw.

10	fetal.tw. or Fetal Development/

11	newborn.tw.

12	1 or 2

13	3 or 4 or 5 or 6 or 7 or 8 or 9 or 10 or 11

14	12 and 13
